# PPAR-delta acts as a metabolic master checkpoint for metastasis in pancreatic cancer

**DOI:** 10.1101/2021.11.15.468579

**Authors:** Beatriz Parejo-Alonso, David Barneda, Sara Trabulo, Sarah Courtois, Sara Compte-Sancerni, Laura Ruiz-Cañas, Quan Zheng, Jiajia Tang, Minchun Chen, Zhenyang Guo, Ulf Schmitz, Pilar Irún, Laure Penin-Peyta, Shanthini M. Crusz, Andres Cano-Galiano, Sergio Lopez-Escalona, Petra Jagust, Pilar Espiau-Romera, Mariia Yuneva, Meng-Lay Lin, Angel Lanas, Bruno Sainz, Christopher Heeschen, Patricia Sancho

**Affiliations:** IIS Aragón, Zaragoza 50009, Spain; Barts Cancer Institute, Queen Mary University of London, United Kingdom; Department of Biochemistry, Autónoma University of Madrid (UAM), School of Medicine, Instituto de Investigaciones Biomédicas (IIBm) “Alberto Sols” CSIC-UAM, 28029 Madrid, Spain; Chronic Diseases and Cancer, Area 3, Instituto Ramón y Cajal de Investigación Sanitaria (IRYCIS), Madrid, Spain; Center for Single-Cell Omics and Key Laboratory of Oncogenes and Related Genes, Shanghai Jiao Tong University School of Medicine, China; Department of Molecular & Cell Biology, College of Public Health, Medical and Veterinary Sciences, James Cook University, Australia; Centro de Investigación Biomédica en Red de Enfermedades Hepáticas y Digestivas (CIBEREHD), Instituto de Salud Carlos III (ISCIII), Zaragoza, Spain; Oncogenes and Tumour Metabolism Laboratory, Francis Crick Institute, 1 Midland Rd, Kings Cross, London NW1 1AT, United Kingdom; Department of Gastroenterology, Hospital Universitario Lozano Blesa. IIS Aragón, Zaragoza 50009, Spain; Pancreatic Cancer Heterogeneity, Candiolo Cancer Institute, FPO-IRCCS, Candiolo, Turin, Italy

**Author notes:** ***Correspondence*:** Dr. Patricia Sancho, PhD, IIS Aragón, Hospital Universitario Miguel Servet, Zaragoza, Spain,; Dr. Christopher Heeschen, MD, PhD, (Center for Single-Cell Omics, Shanghai Jiao Tong University School of Medicine, China.

**Keywords:** Pancreatic ductal adenocarcinoma, Epithelial-to-mesenchymal transition, Metastasis, Cancer stem cells, Metabolism, MYC, PGC1A, PPAR, PPARD

## Abstract

In pancreatic cancer, emerging evidence suggests that PPAR-δ overexpression is associated with tumor progression and metastasis, but a mechanistic link is still missing. Here we now show that PPAR-δ acts as the integrating upstream regulator for the metabolic rewiring, which is preceding the subsequent initiation of an invasive/metastatic program. Specifically, paracrine and metabolic cues regularly found in the metastasis-promoting tumor stroma consistently enhance, via induction of PPAR-δ activity, the glycolytic capacity and reserve of pancreatic cancer cells, respectively, accompanied by decreased mitochondrial oxygen consumption. Consequently, genetic or pharmacological inhibition of PPAR-δ results in reduced invasiveness and metastasis. Mechanistically, PPAR-δ acts by shifting the *MYC/PGC1A* balance towards *MY*C, enhancing metabolic plasticity. Targeting *MYC* similarly prevents the metabolic switch and subsequent initiation of invasiveness. Therefore, our data demonstrate that PPAR-δ is a key initiator for the metabolic reprogramming in pancreatic cancer, thereby acting as a checkpoint for the phenotypic change towards invasiveness. These findings provide compelling evidence for a novel treatment strategy to combat pancreatic cancer progression.

## Introduction

Pancreatic Ductal Adenocarcinoma (PDAC), the most frequent form of pancreatic cancer is an extremely lethal disease with high metastatic potential (Hidalgo, 2010). At the time of diagnosis, 80-90% of the patients are already at an advanced/metastatic disease stage, with very limited therapeutic options and a particularly poor long-term outcome (Siegel et al., 2017). This can, at least in part, be attributed to the hierarchical organization of PDAC, containing cells with tumor-initiating properties or cancer stem cells (CSCs), which constitute the driving force for disease progression, metastasis, and chemo-resistance (Hermann et al., 2007; Li et al., 2007).

CSCs are capable of unlimited self-renewal, thereby maintaining the CSC pool and also giving rise to the more differentiated progenies (non-CSCs) with a high proliferative capacity. Although both CSCs and non-CSCs can acquire mobility by processes such as epithelial-to-mesenchymal transition (EMT), the arising metastatic CSCs would predominantly be able to initiate secondary lesions due to their strong tumor-initiating capacity. Thus, complementing current chemotherapies with strategies that efficiently target CSCs, bears the potential to eventually improve patients’ long-term survival (Gallmeier et al., 2011; Lonardo et al., 2011; Mueller et al., 2009; Zhang et al., 2016).

We recently reported that *c-MYC* (hereinafter referred to as *MYC*) plays an essential role in defining the metabolic phenotype and stemness of PDAC cells, by negatively controlling the expression of the mitochondrial biogenesis factor *PGC1A* (Peroxisome proliferator-activated receptor gamma coactivator 1-alpha) (Sancho et al., 2015). Reduced *MYC* expression in CSCs was required to unleash *PGC1A* and promoted an OXPHOS-dependent metabolic phenotype, thereby enhancing their self-renewal capacity. This rendered CSCs particularly sensitive to mitochondrial targeting (i.e. Metformin), whereas differentiated cancer cells, characterized by increased *MYC* expression and a glycolytic phenotype, were not sensitive to Metformin.

Intriguingly, however, a subpopulation of CSCs turned out to be resistant to mitochondrial inhibition due to an increased *MYC/PGC1A* ratio and metabolic plasticity, allowing them to modulate their metabolism in response to exogenous environmental cues. This subset of Metformin-resistant CSCs displayed a highly invasive phenotype, suggesting a potential link between metabolism and invasiveness. Indeed, here we now conclusively show that metabolic reprogramming induced by PPAR-δ (Peroxisome Proliferator-Activated Receptor delta) via enhancing *MYC/PGC1A* ratio, which precedes and facilitates the acquisition of an invasive, EMT-like phenotype in PDAC cancer (stem) cells. This phenotype was induced either through partial inhibition of mitochondrial activity and nutrient stress, respectively, or via stromal cues. Single-cell RNAseq identified PPAR-δ as a directly druggable upstream target, which integrates both nutrient-sensing and stromal signals to modulate cellular metabolism and subsequently invasiveness and metastasis via increasing the *MYC/PGC1A* ratio. Therefore, targeting PPAR-δ represents a novel and translatable approach to counteract PDAC progression and metastasis.

## Results

### Induction of an EMT-like phenotype in PDAC cells

We have previously shown while prolonged treatment of PDAC cultures with the mitochondrial complex I inhibitor Metformin eliminated a large fraction of CSCs, outgrowth of pre-existing resistant CSC clones occurred (Sancho et al., 2015; Lonardo et al., 2013). These prevalent Metformin-resistant cells were morphologically distinct with an elongated shape and diminished cell-to-cell contact and showed upregulation of EMT-related genes, e.g. *VIM* and *ZEB1* (**Figure S1A**).

To determine if the acquisition of an EMT-like phenotype could be a general downstream consequence of diminished mitochondrial activity, we next treated various primary PDAC cultures using distinct means to inhibit their mitochondrial functions, e.g. reducing mitochondrial uptake of different carbon sources or diminishing the activity of the electron transport chain (ETC). Indeed, short-term treatment with Malonate (complex II inhibitor), Etomoxir (mitochondrial long-chain fatty acid transporter blocker) and UK5099 (mitochondrial pyruvate carrier blocker) resulted in morphological and gene expression changes in the cells that are consistent with the induction of EMT (**Figure 1A, S1B**). Interestingly, mimicking conditions frequently found in the tumor microenvironment (low pH, nutrient deprivation, hypoxia) induced similar alterations in morphology and gene expression (**Figure 1A, S1B**). Even glucose or glutamine deprivation alone induced expression of EMT genes (**Figure S1C**). Thus, decreased mitochondrial activity, either directly induced by inhibitors or indirectly by diminishing metabolic substrates, consistently led to the induction of an EMT-like phenotype.

**Figure 1.**
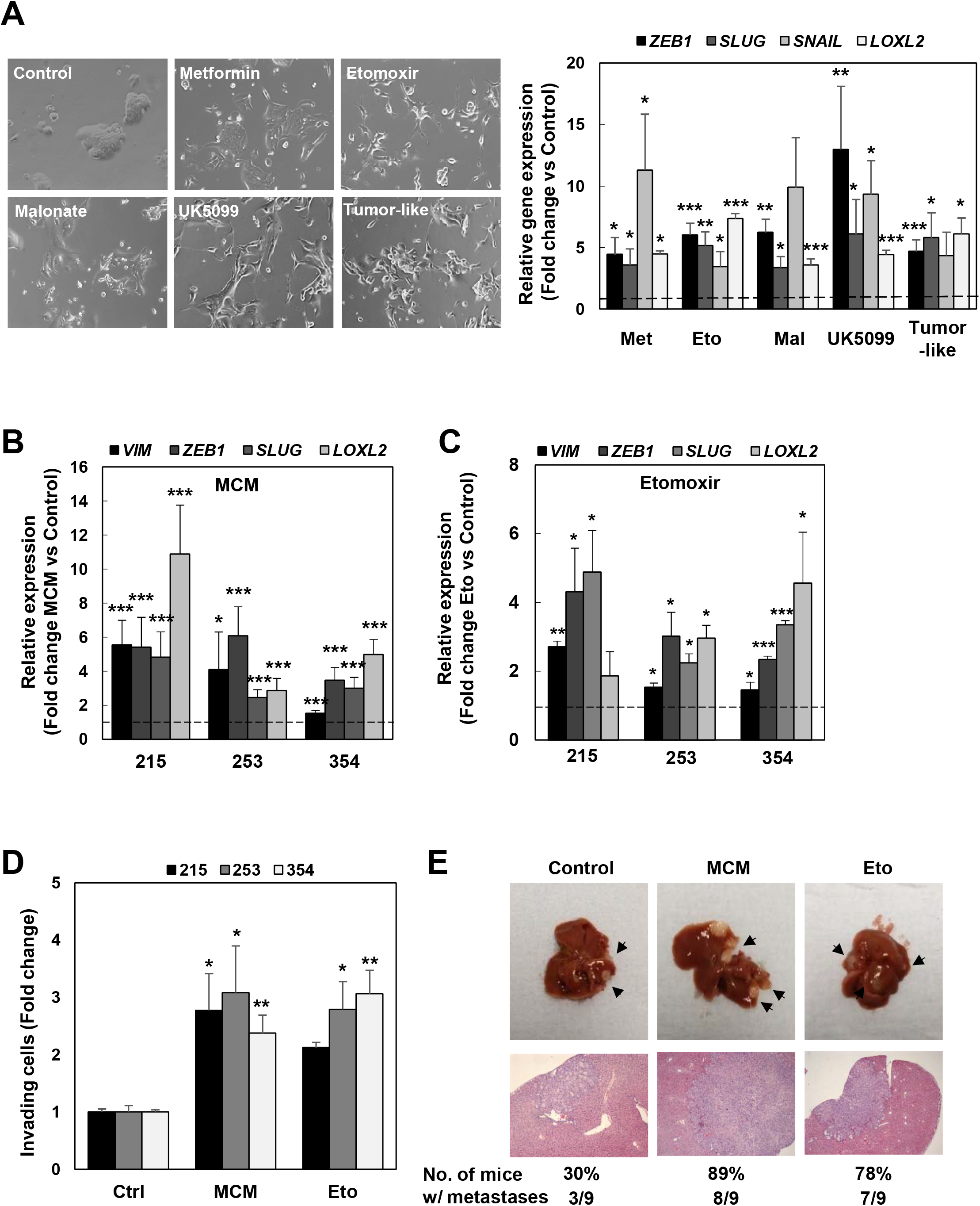
Induction of EMT-like phenotype in PDAC cells. (**A**) **Left**: Representative images illustrating morphological changes for PDAC-354 cells in response to treatment for 72h with the complex I inhibitor Metformin (3mM), the ß-oxidation inhibitor Etomoxir (20µM), complex II inhibitor Malonate (5mM), the pyruvate carrier inhibitor UK5099 (100µM), or tumor-like conditions (low pH (HCl 50 μM) + low glucose concentration (1 mM) + 3% O_2_). **Right**: Expression of EMT-associated genes (*VIM* [vimentin], *SNAIL, SLUG, ZEB1* and *LOXL2*) was determined by rtQPCR after cells were treated for 48h as indicated. Pooled data for PDAC-185, A6L, 215, 253, and 354 (n≤4 for each cell type). Data are normalized to *HPRT* (**lower left panel**). (**B-C**) PDAC-215, 253, and 354 cells were treated with macrophage-conditioned medium (MCM) or 20µM Etomoxir (Eto) for 48h and expression of EMT-associated genes (*VIM, SNAIL, ZEB1, SLUG* and *LOXL2*) was determined by rtQPCR (n≤4 for each cell type). Data are normalized to *HPRT*. (**D**) Cells were treated as indicated above and seeded in modified Boyden invasion chambers containing 20% FBS in the lower compartment. The number of invasive cells was analyzed after 16h. (**E**) GFP^+^ Luciferase^+^ PDAC-354 cells were treated with control, MCM, or 20µM Eto for 48h and then injected intrasplenically to assess their metastatic capacity. Representative photographs of liver metastasis and subsequent H&E staining. All data are represented as mean ± SEM. * p<0.05, ** p<0.01, *** p<0.001. See also Figure S1.

We previously identified microenvironmental signals from tumor-associated macrophages (TAMs) and pancreatic stellate cells (PSCs) that strongly induce invasion and metastasis in PDAC (Lonardo et al., 2012; Sainz et al., 2014, 2015). Co-culturing PDAC cells with primary human TAMs or PSCs resulted in up-regulation of *VIM* and *ZEB1* (**Figure S1D**). The observed changes in gene expression induced by TAMs could be mimicked by macrophage-conditioned medium (MCM) (**Figure 1B)**, and were comparable to the changes induced by Etomoxir **(Figure 1C)**. Treatment with MCM or Etomoxir consistently upregulated *ZEB1* in both CD133^+^ CSCs and CD133^−^ non-CSCs, independent of their mitochondrial content (**Figure S1E**), but did not significantly alter their self-renewal capacity (**Figure S1F**). In line with the outlined morphological and transcriptional changes, the cells in both models showed a consistent and strong induction of *in vitro* invasiveness and *in vivo* metastasis, respectively (**Figure 1D, 1E**).

From this diverse panel of invasion/metastasis-inducers, we selected MCM and Etomoxir as the most suitable and relevant stimuli for our subsequent studies. This selection was based on their distinct mechanism of EMT induction: 1) microenvironmental signals from TAMs (MCM) and 2) partial impairment of mitochondrial activity by Etomoxir-mediated inhibition of fatty acid uptake, which resulted in a comparable and reproducible increase of cell invasiveness *in vitro* and metastasis *in vivo* (**Figure 1D, 1E**).

### A common transcriptional program linked to PPARD controls invasiveness and metastasis induced by microenvironmental signals

In order to detect common global transcriptional changes induced by both MCM and Etomoxir, we next performed single-cell RNAseq (scRNAseq) analyses for three different PDAC models. Notably, scRNAseq showed that the majority of cells underwent a strong induction of the Hallmark EMT signature, whereas a smaller subset of cells did not respond to the EMT cues (e.g. Cluster 2, **Figure 2A**). As expected from their distinct mechanism of action, distinct transcriptional profiles in response to EMT induction were noted for MCM and Etomoxir (**Figure 2SA**). These findings were consistent with the diverse morphological changes upon induction of EMT where a subset of cancer cells maintained their epithelial morphology (**Figure S2B**).

**Figure 2.**
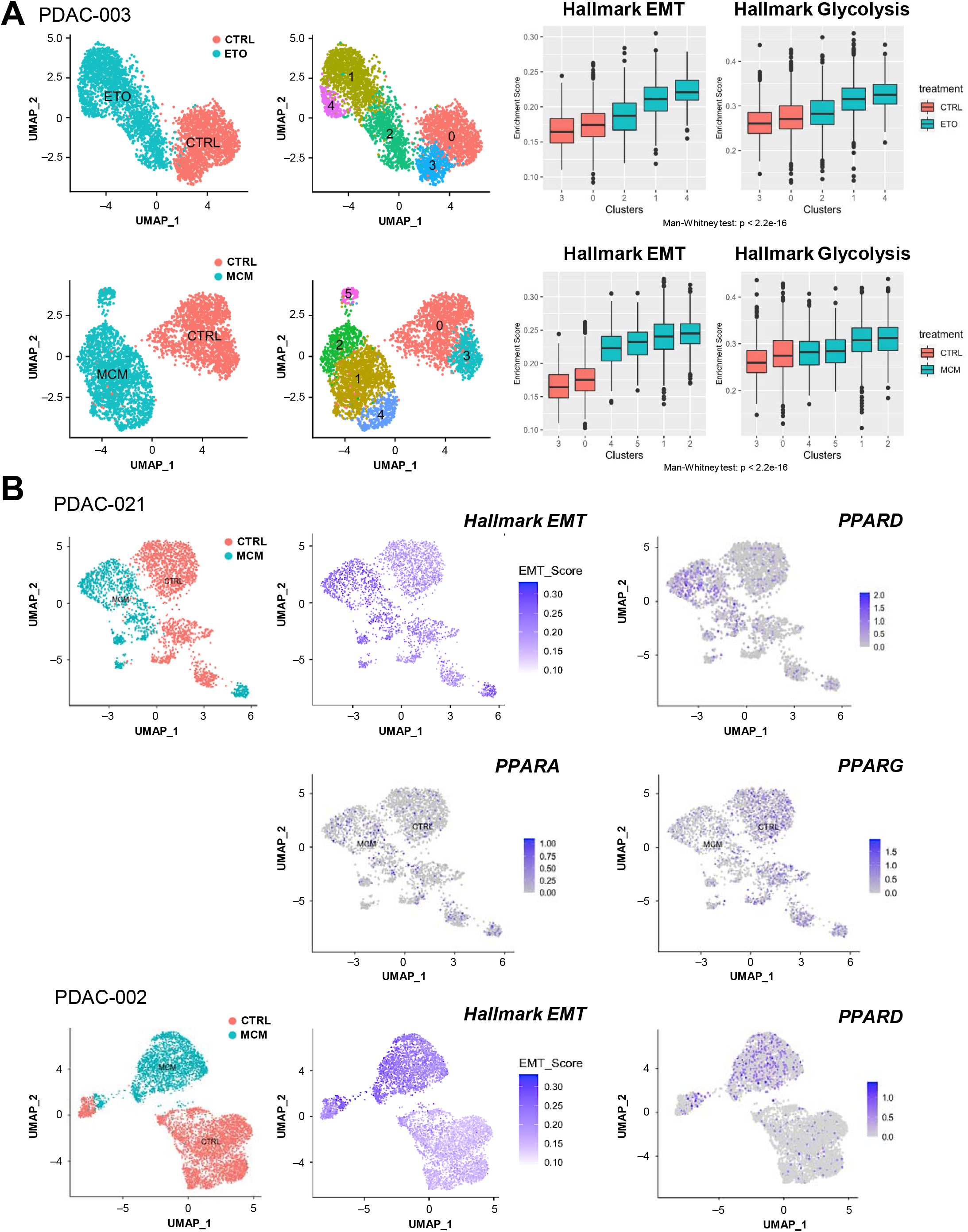
Single-cell RNAseq analysis identifies metabolic switch during EMT induction. (**A**) **Left panel**: PDAC-003 cells were treated with control vehicle (CTRL), macrophage-conditioned medium (MCM), or 20µM Etomoxir (ETO) for 48h to induce an EMT-like state and were then subjected to single-cell RNAseq (10X Genomics Chromium platform). Unsupervised clustering of viable PDAC cells exposed to CTRL, MCM or ETO, represented as UMAP plots. Different clusters are color coded. **Right panel**: Boxplots illustrating gene set enrichment results for the *EMT* and *Glycolysis* (Hallmark data set) for different clusters in CTRL versus MCM and ETO treatment, respectively. Differences in enrichment scores between treatments were assessed using the Mann-Whitney U test. (**B**) Expression of EMT hallmark signature and PPARD family members in single cancer cells (PDAC-002 and 021) displayed as unsupervised clusters and color-coded for allocated treatment.

Intriguingly, while MCM and Etomoxir induced a distinct transcriptional profile compared to untreated control cells (**Figure S2A**), Gene Set Enrichment Analysis (GSEA) analysis revealed that both treatments consistently activated metabolic pathways such as glycolysis and hypoxia, an effect that again was mostly confined to cells responding to EMT induction (**Figure 2A, S2C**). Bulk transcriptional analysis showed a similar trend, although differences were less pronounced, most likely due to contained cells that did not respond to EMT induction (**Figure S2C, S2D**). Together, these data demonstrate that the majority of PDAC cells undergo similar metabolic changes in response to EMT induction.

We then further analyzed the scRNAseq data sets to identify specific metabolism-related genes and regulators. Most intriguingly, upon induction of EMT we noted a consistent upregulation of the nuclear Peroxisome Proliferator-activated Receptor-δ (*PPARD*) across the different clusters (**Figure 2B**). While *PPARD* upregulation was heterogeneous, it was mostly confined to cells displaying the Hallmark EMT signature. *PPARD* is a member of the *PPAR* subfamily of nuclear hormone receptors, together with *PPARA* and *PPARG*. This subfamily modulates energy homeostasis by controlling the expression of numerous genes involved in lipid and glucose metabolism (Dubois et al., 2017). Notable, we only found *PPARD* to be consistently upregulated in EMT cells, whereas the expression of other family members, e.g. *PPARA* and *PPARG*, was not altered (**Figure 2B**).

We next performed a series of bioinformatic analyses of publicly accessible human datasets, to further interrogate a possible association of these nuclear receptors with human PDAC aggressiveness and metastasis. First, analysis of TCGA and GTEx datasets (http://gepia.cancer-pku.cn/index.html) showed significantly increased expression levels for the *PPAR* family members *PPARD* and *PPARG* for tumor tissue versus normal tissue (**Figure 3A**), which also correlated with poor outcome (**Figure 3B**). Interestingly, only *PPARD* expression positively correlated with an EMT-related gene signature formed by *ZEB1, SNAIL* and *SLUG* in the tumor (**Figure 3C**). We performed GSEA of the TCGA dataset and compared samples belonging to the top and bottom quartiles of *PPARD* expression. Applying the Hallmark gene set collection, we found that the EMT pathway was one of the most significantly enriched pathways in patients with high *PPARD* expression, together with metabolism-related pathways glycolysis and hypoxia (**Figure 3D, 3E**). Consistently, the OXPHOS pathway was significantly downregulated in the high *PPARD* expression quartile (**Figure 3D, 3E**). Together, these results mirror the transcriptional expression pattern induced by our *in vitro* EMT conditions, further corroborating how hypothesis that *PPARD* acts as a key regulator for the metastatic program in PDAC.

**Figure 3.**
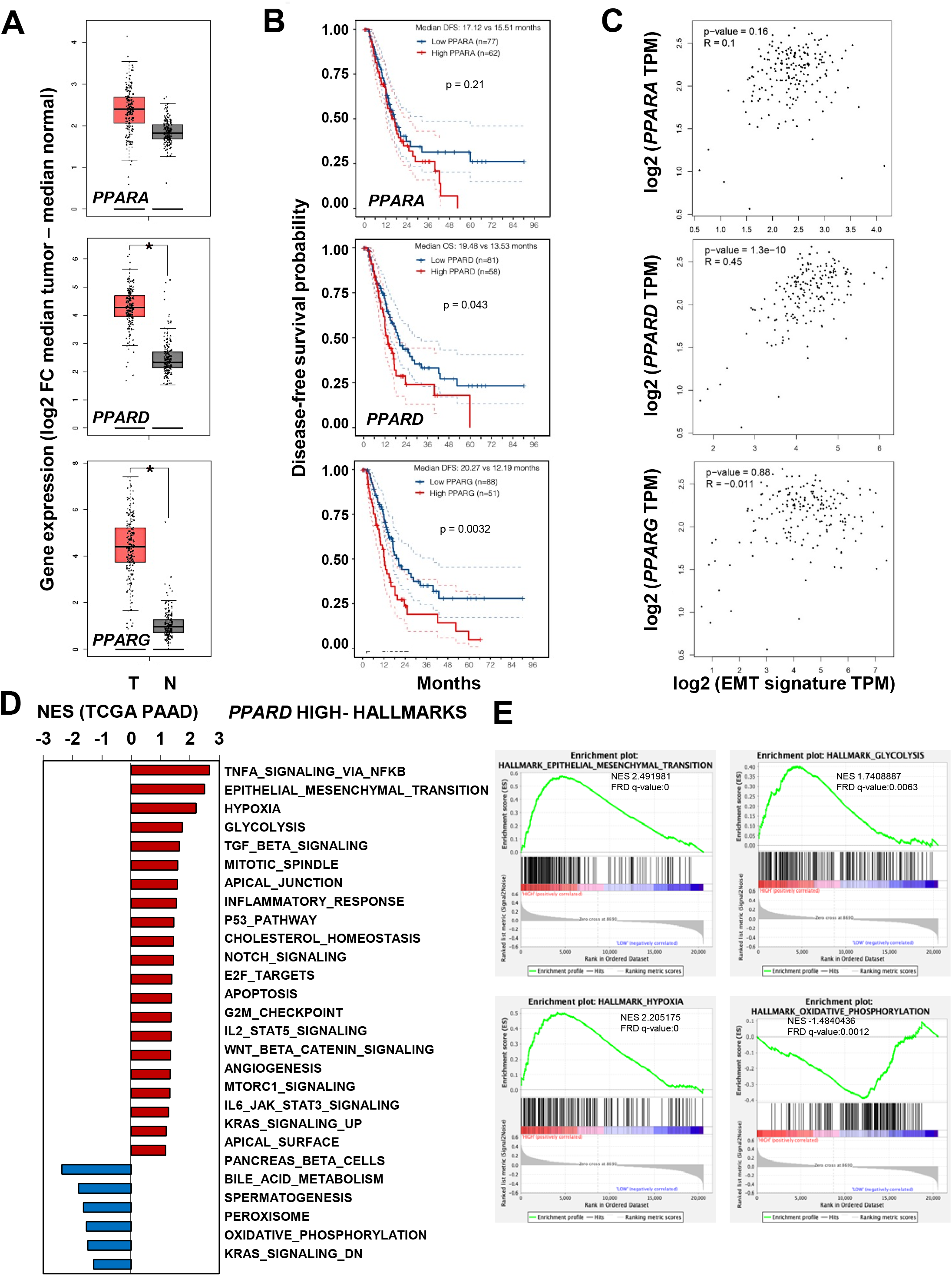
*PPARD* expression is linked to metabolic switch and invasiveness in PDAC patients. (**A**) Expression levels for PPAR family members in PDAC tumors (T) versus surrounding normal tissue (N) included in the TCGA and GTEx projects. (**B**) Patients were dichotomized for the tumor expression levels for PPAR family members (higher and lower expression compared to the mean; RNA Seq V2 RSEM values). Kaplan Meier survival curves for disease-free survival are shown. Dotted lines denote the confidence interval. (**C**) Correlation between tumor expression levels for PPAR family members and an EMT-associated signature composed of *SLUG, SNAIL*, and *ZEB1*. (**D**) Gene sets enriched in the transcriptional profile of tumors belonging to the top *PPARD* high-expression group, compared with the bottom expression group in the TCGA data series. Shown are the NES (normalized enrichment score) values for each pathway using the Hallmark gene sets, meeting the significance criteria: nominal p-value of <0.05, FDR□< □25%. (**E**) Enrichment plot for EMT, Glycolysis, Hypoxia and OXPHOS hallmarks in *PPARD* high versus low samples, showing values of NES and FRD q-values.

### PPAR-δ controls invasiveness and metastasis in PDAC

Using our panel of five inducers of EMT, we were able to confirm a consistent upregulation of *PPARD*, irrespective of the trigger (**Figure 4A**). Although MCM and Etomoxir also upregulated other members of the *PPAR* family, i.e. *PPARA* and *PPARG* (**Figure S3A, S3B**), we found that only *PPARD* was significantly up-regulated during the first 24h, when changes of cellular morphology and *ZEB1* expression were still minor or even undetectable (**Figure S3A, S3B**). The exclusive and rapid PPAR-δ activation within 24h could be further corroborated by demonstrating direct binding to its consensus sequence (**Figure 4B**) and preferential up-regulation of PPAR-δ target genes (**Figure S3C**).

**Figure 4.**
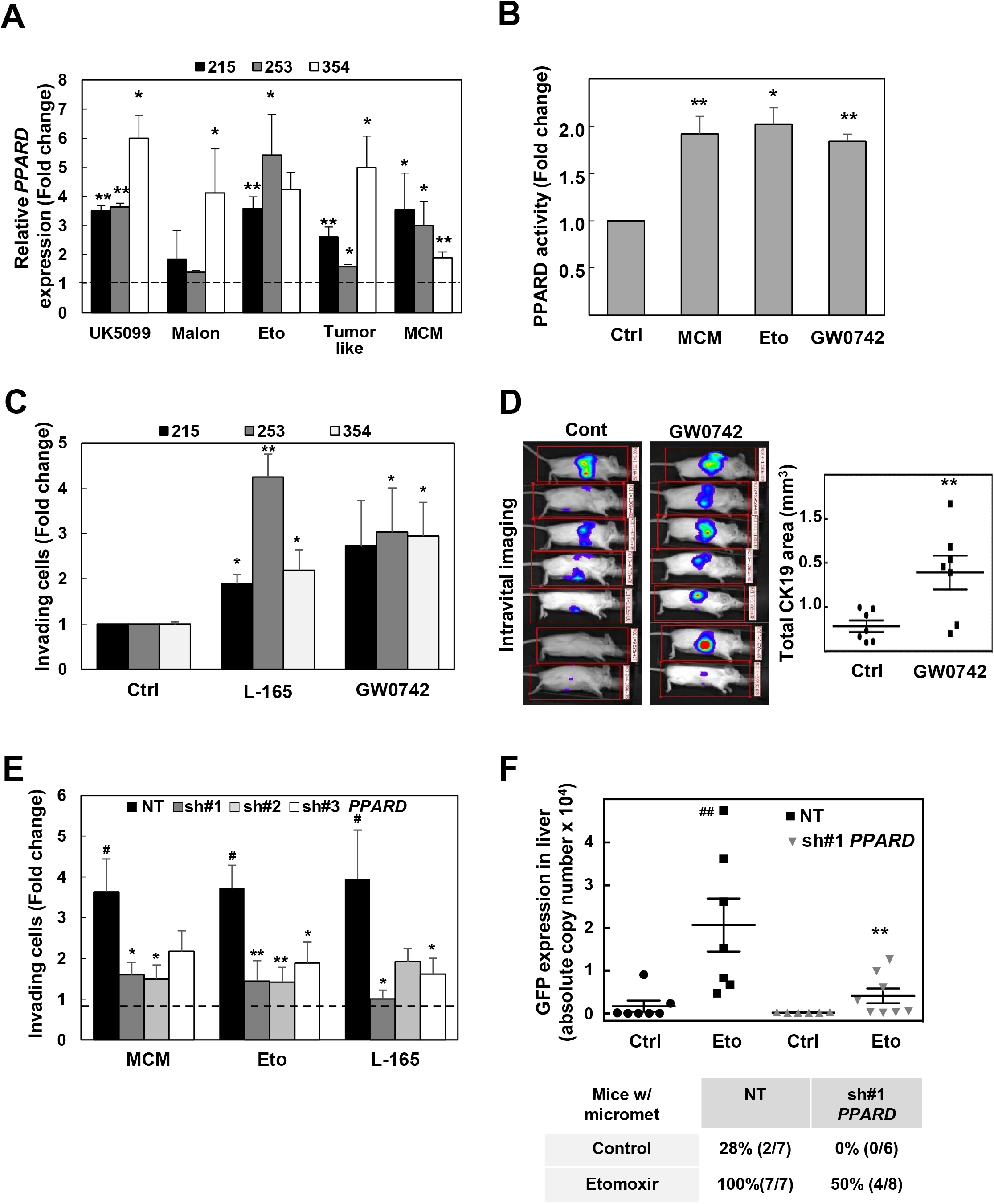
Activation of PPAR-δ initiates invasiveness and metastasis. **(A)** *PPARD* expression upon 48h of treatment with the complex I inhibitor Metformin (3mM), the ß-oxidation inhibitor Etomoxir (20µM), complex II inhibitor Malonate (5mM), the pyruvate carrier inhibitor UK5099 (100µM), or tumor-like conditions (HCl 50 µM + 1 mM Glc+ 3% O2) with the indicated stimuli in PDAC-215, 253, and 354 cells. (**B)** PPAR-δ activity, measured as binding to its specific DNA sequence, following stimulation with MCM, Eto, and PPAR-δ agonist GW0742 for 24 hours. (**C**) Invasive capacity of cells treated for 48h with the PPAR-δ agonists L-165 and GW0742, respectively. Cells were placed in modified Boyden invasion chambers containing 20% FBS in the lower compartment and the number of invasive cells was assessed after 16h. (**D**) *In vivo* metastatic activity of PDAC-354-GFP-luc cells pretreated with GW0742 for 48h. After surgery, mice received three more daily doses of GW0742 (0.3mg/kg i.v.). IVIS imaging (**left panel**) and quantification of the total CK19 area in the livers 9 weeks after implantation (**right panel**). (**E**) PDAC-215, 253, and 354 cells were stably transduced with inducible lentiviral vectors expressing either a non-targeting shRNA (NT) or three different shRNAs against PPARD (sh#1, sh#2, sh#3). Transduced cells were pre-treated with doxycycline for 24h, then incubated with MCM, Eto, or L-165 for 48h. (**F**) ZsGreen expression by rt-QPCR in liver homogenates from an *in vivo* metastasis assay of PDAC-354 cells stably expressing either the NT or the sh#1 against PPARD. Cells were pretreated with doxycycline and/or 20 Eto μM for 48h. After intrasplenic implantation, mice were treated with oral doxycycline (2mg/ml drinking water) and Etomoxir (15 mg/kg, i.p. every day) for 7 days, when splenectomies were performed. Table indicates the percentage and total number of micrometastases in each experimental group. All data are represented as mean ± SEM. * p<0.05, ** p<0.01, *** p<0.001. See also Figures S3 and S4.

Importantly, treatment of PDAC cells with PPAR-δ chemical agonists (GW0742, GW501516, and L-165), but not PPAR-α or PPAR-γ agonists (e.g. WY14643 and rosiglitazone), resulted in a dose-dependent induction of EMT-related genes and typical morphological changes (**Figure S4A, S4B**). Conversely, knockdown of *PPARD* (**Figure S4C**) virtually abrogated the transcriptional changes induced by MCM, Etomoxir and the PPAR-δ agonist L-165 (**Figure S4D**). Functionally, PPAR-δ activation by agonists resulted in enhanced invasiveness *in vitro* (**Figure 4C**) and metastasis *in vivo* (**Figure 4D**). Moreover, knockdown of *PPARD* reversed MCM, Etomoxir, or PPAR-δ agonist-induced invasiveness (**Figure 4E**) as well as Etomoxir-induced metastasis *in vivo* (**Figure 4F**). Together, these data demonstrate that PPAR-δ, but not other PPARs, is responsible for transcriptional and functional changes concomitant with EMT induction, thereby strongly suggesting an essential role for PPAR-δ in the process of cancer cell invasiveness and metastasis.

### PPAR-δ controls a metabolic program linked to invasiveness and metastasis in PDAC

As shown above, EMT induction by microenvironmental signals was strongly linked to a metabolic transcriptional program characterized by glycolysis and hypoxia signaling induction and OXPHOS inhibition (**Figure 2**); features also observed in patients expressing high *PPARD* levels (**Figure 3D, 3E**). Interestingly, using a carbohydrate metabolism PCR array, we found genes implicated in uptake and intermediary metabolism of alternative sugars such as fructose, TCA substrates, amino acids and lipids to be commonly upregulated following EMT induction with MCM, Etomoxir, or the pyruvate carrier inhibitor UK5099 (**Figure 5A**).

**Figure 5.**
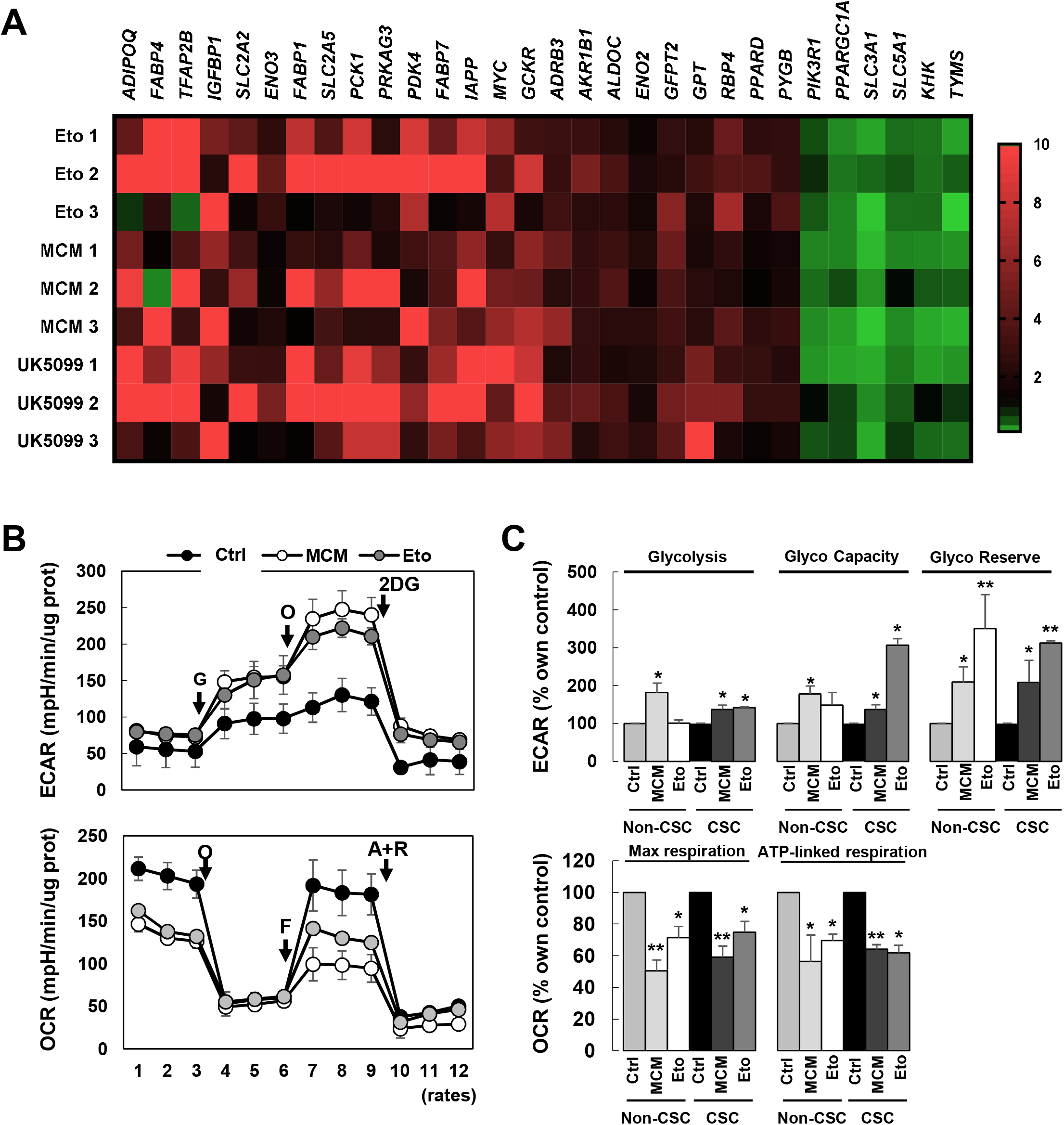
A common PPAR-δ-initiated metabolic program drives invasiveness. (**A**) Gene expression profile as assessed by a Carbohydrate metabolism PCR array in PDAC-354 cells. Heatmap showing only genes whose expression was significantly altered. Cells were treated with vehicle (Cont), macrophage-conditioned medium (MCM), 20µM Etomoxir (Eto), or 100µM of the pyruvate carrier inhibitor UK5099 for 48h. (**B)** Representative Extracellular Acidification Rate (ECAR) profile for PDAC-253 cells (Glycolysis test) (**upper panel**). G, Glucose; O, ATP synthase inhibitor Oligomycin; 2DG, Glycolysis inhibitor 2-deoxy-glucose. Representative Oxygen Consumption Rate (OCR) profile for PDAC-253 cells (Mitochondrial stress test) (**lower panel**). O, ATP synthase inhibitor Oligomycin; F, mitochondrial oxidative phosphorylation uncoupler FCCP (Carbonyl cyanide-4 [trifluoromethoxy] phenylhydrazone); A+R, complex III inhibitor Antimycin A + Electron transport change inhibitor Rotenone. (**C**) Glycolysis, glycolytic capacity, and reserve in adherent vs sphere-derived cells (**upper panel**). Pooled data from PDAC-215, 253, and 354. Maximal and ATP-linked respiration in non-CSCs vs CSCs (**lower panel**). Pooled data for PDAC-215, 253, and 354. All data are represented as mean ± SEM. * p<0.05, ** p<0.01, *** p<0.001. See also Figure S3.

Moreover, the PCR array confirmed a significant increase of *PPARD* and a switch in the *MYC/PGC1A* balance towards increased *MYC* expression (**Figure 5A**). We had previously described a similar switch in Metformin-resistant primary PDAC cells (Sancho et al., 2015), favoring glucose metabolism via glycolysis versus OXPHOS. As predicted by the above transcriptional profiling, metabolic parameters associated with enhanced glycolytic activity (glycolysis, glycolytic capacity and reserve) were increased upon induction of EMT with MCM or Etomoxir in CSCs and non-CSCs (**Figure 5B, 5C**). Conversely, mitochondrial oxygen consumption rate (OCR) was reduced upon pretreatment with MCM or Etomoxir. Both maximal and ATP-linked OCR were inhibited by 40-50% upon the indicated treatments, with similar changes in CSCs and non-CSCs (**Figure 5B, 5C**), despite their different baseline levels (Sancho et al., 2015). These effects on OCR could be mimicked by co-culture of the cancer cells with primary human TAMs or PSCs (**Figure S5A-C**).

Of note, metabolic changes related to glycolysis were less evident (**Figure 5C**), corroborated by a slight enhancement of glucose uptake and release of lactate and alanine upon treatment (**Figure S5D-F**). However, both glycolytic capacity and reserve, which measure metabolic plasticity as the ability to switch to alternative pathways upon complete inhibition of mitochondrial ATP, were increased upon EMT induction (**Figure 5C**). We therefore hypothesized that EMT induction favors the combined metabolism of glucose by glycolysis together with the use of alternative carbon sources in mitochondria, as suggested by the PCR array data. This would be particularly relevant for CSC functionality as most of these cells in the native state lack metabolic plasticity and are unable to compensate mitochondrial impairment by switching to glycolysis (Sancho et al., 2015).

To further test this hypothesis, we performed a series of experiments manipulating *PPARD* expression and function by genetic and pharmacological means. Functionally, when PPAR-δ induction was prevented by inducible knockdown, the metabolic changes associated with EMT, e.g. increased glycolytic capacity and diminished mitochondrial respiration, were abrogated (**Figure 6A**). Conversely, treatment of PDAC cells with the chemical agonists GW0742 and L-165, which specifically activated and upregulated *PPARD* (**Figure 4B, S3D**), recapitulated the metabolic switch induced by induction of EMT (**Figure 6B**). Consistent with PPARs stimulation of lipid metabolism and, specifically, fatty acid oxidation (FAO), decreased glucose-dependent respiration was completely rescued by the addition of palmitate and carnitine to the culture medium (**Figure 6C**). This suggests that PPAR-δ promoted glucose diversion to glycolysis while upregulating the FAO machinery to provide an alternative carbon source for TCA cycle when substrates are available.

**Figure 6.**
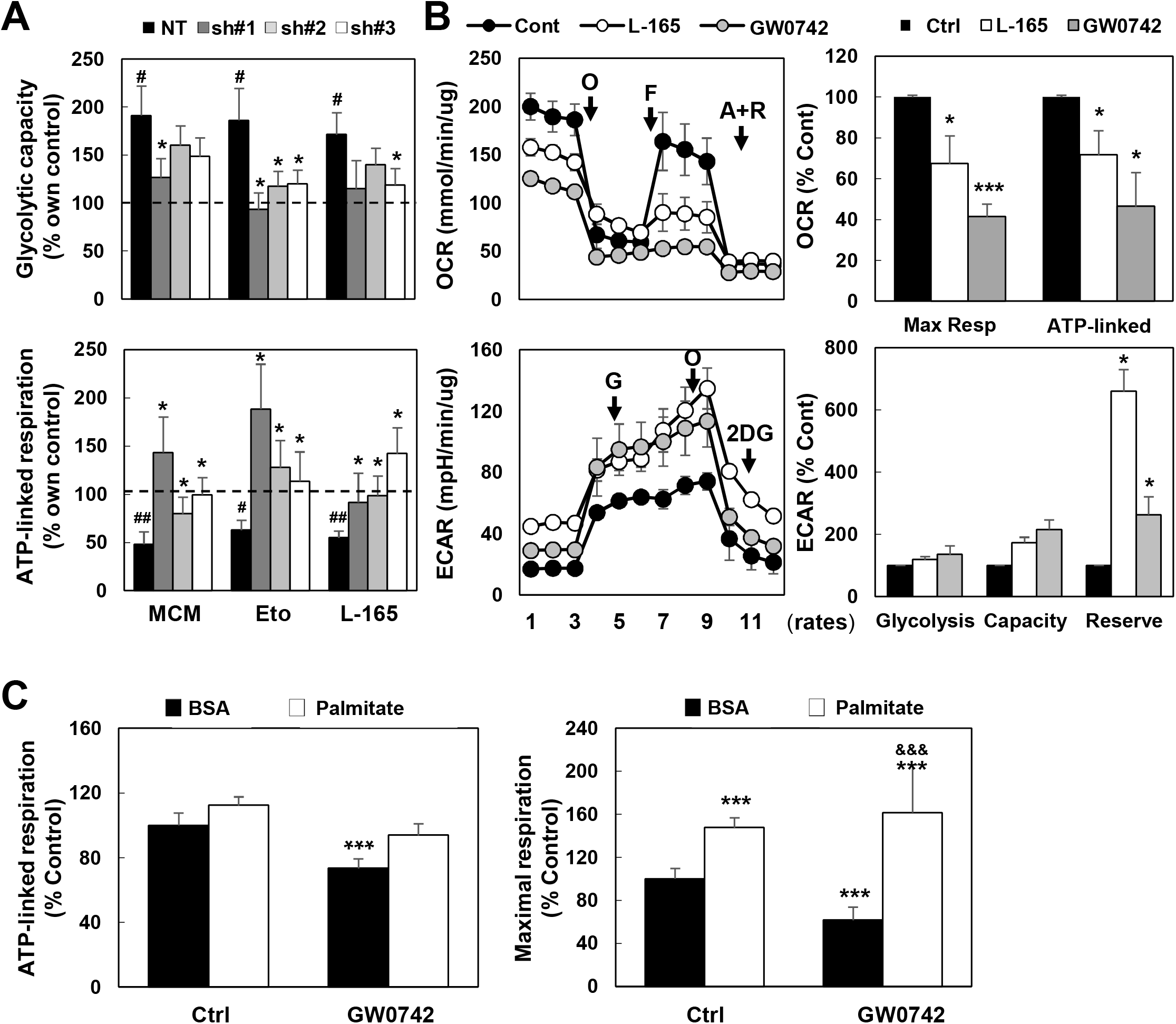
PPAR-δ controls the balance between OXPHOS and glycolysis, linked to EMT and metastasis. (**A**) PDAC-215, 253, and 354 cells were stably transduced with inducible lentiviral vectors expressing either a non-targeting shRNA (NT) or three different shRNAs against PPARD (sh#1, sh#2, sh#3). Transduced cells were pre-treated with doxycycline for 24h, then incubated with macrophage-conditioned medium (MCM), Etomoxir (Eto), or L-165 and tested for glycolytic capacity (**upper panel**) and ATP-linked respiration (**lower panel**) after additional 24h. (**B**) Mitochondrial stress test (**upper row**) and glycolysis test (**lower row**) following treatment with control (Cont) or the PPAR-δ agonists L-165 or GW0742. **Left column**, representative OCR and ECAR profiles for PDAC-253. **Right column**, pooled data for PDX-215, 253, and 354 cells. O, ATP synthase inhibitor Oligomycin; F, mitochondrial oxidative phosphorylation uncoupler FCCP (Carbonyl cyanide-4 [trifluoromethoxy] phenylhydrazone); A+R, complex III inhibitor Antimycin A + Electron transport change inhibitor Rotenone. G, Glucose; 2DG, Glycolysis inhibitor 2-deoxy-glucose. * p<0.05, ** p<0.01. (**C**) ATP-linked respiration (**left panel**) and maximal respiration (**right panel**) for control versus GW0742-treated cells following treatment with or without Palmitate-BSA (FAO assay). Cells were treated with 10 μM GW0742 for 48 hours prior to the assay. Pooled data from PDAC-215, 253 and 354 cells. All data are represented as mean ± SEM. *** p<0.001 vs Control, &&& p<0.001 vs Palmitate.

### Downstream signaling cascade initiating the metabolic switch and promoting invasiveness

*MYC* plays an essential role in defining the metabolic phenotype and stemness of PDAC cells by negatively controlling the expression of the mitochondrial biogenesis factor PGC-1α (Sancho et al., 2015). On the other hand, decreased PGC-1α expression was shown to be essential for inducing migration and metastasis in melanoma and prostate cancer (Luo et al., 2016; Torrano et al., 2016). Here, in addition to the changes in *MYC* and *PGC1A* expression upon treatment with MCM, Etomoxir, and UK5099 (**Figure 5A**), we found that direct overexpression of *MYC* induced an EMT-like phenotype (**Figure S6A**). Together, these data suggest an intricate link between the altered *MYC/PGC1A* balance and the subsequent induction of invasiveness/metastasis. This metabolic reprogramming could be either cause or consequence of acquiring a migratory/metastatic phenotype.

We hypothesized that the PPAR-δ-mediated induction of metastatic activity was related to changes in the *MYC/PGC1A* ratio. To test this hypothesis, we first analyzed the *MYC/PGC1A* ratio in our diverse EMT/metastasis models. Indeed, induction of metastatic activity by diminishing mitochondrial activity due to ETC inhibition or lack of fuel resulted in a consistent increase of the *MYC/PGC1A* ratio (**Figure 7A, S6B**). Notably, the absolute changes in the expression of either MYC or PGC1A individually did not always correlate with the induction of the EMT-like phenotype. For some models, the increase in *MYC* expression was rather modest or absent, whereas *PGC1A* was still greatly reduced and *vice versa*. Instead, we found that, at the mRNA level, an increased *MYC/PGC1A* ratio was most consistently linked to an EMT phenotypic induction. At the protein level, however, MCM, Etomoxir, and the PPAR-δ agonist GW0742 reproducibly increased MYC expression and diminished PGC-1α expression (**Figure 7B**). Moreover, *PPARD* knockdown, which inhibited invasion and metastasis (**Figure 4E-4F**), diminished the increase in the *MYC/PGC1A* ratio induced by MCM, Etomoxir, and PPAR-δ agonist L-165, respectively (**Figure S6C**). Finally, *PPARD* overexpression or PPAR-δ agonist treatment consistently induced *MYC* promoter activity and subsequently reduced *PGC1A* promoter activity (**Figure 7C**), indicating a direct link between *PPARD* expression and the *MYC/PGC1A* ratio. Indeed, the enhanced invasiveness of the cancer cells following treatment with the PPAR-δ agonist could be reversed by either *MYC* knockdown or *PGC1A* overexpression (**Figure 7D**), essentially attributing PPAR-δ’s pro-metastatic effects to its ability to increase the *MYC/PGC1A* ratio.

**Figure 7.**
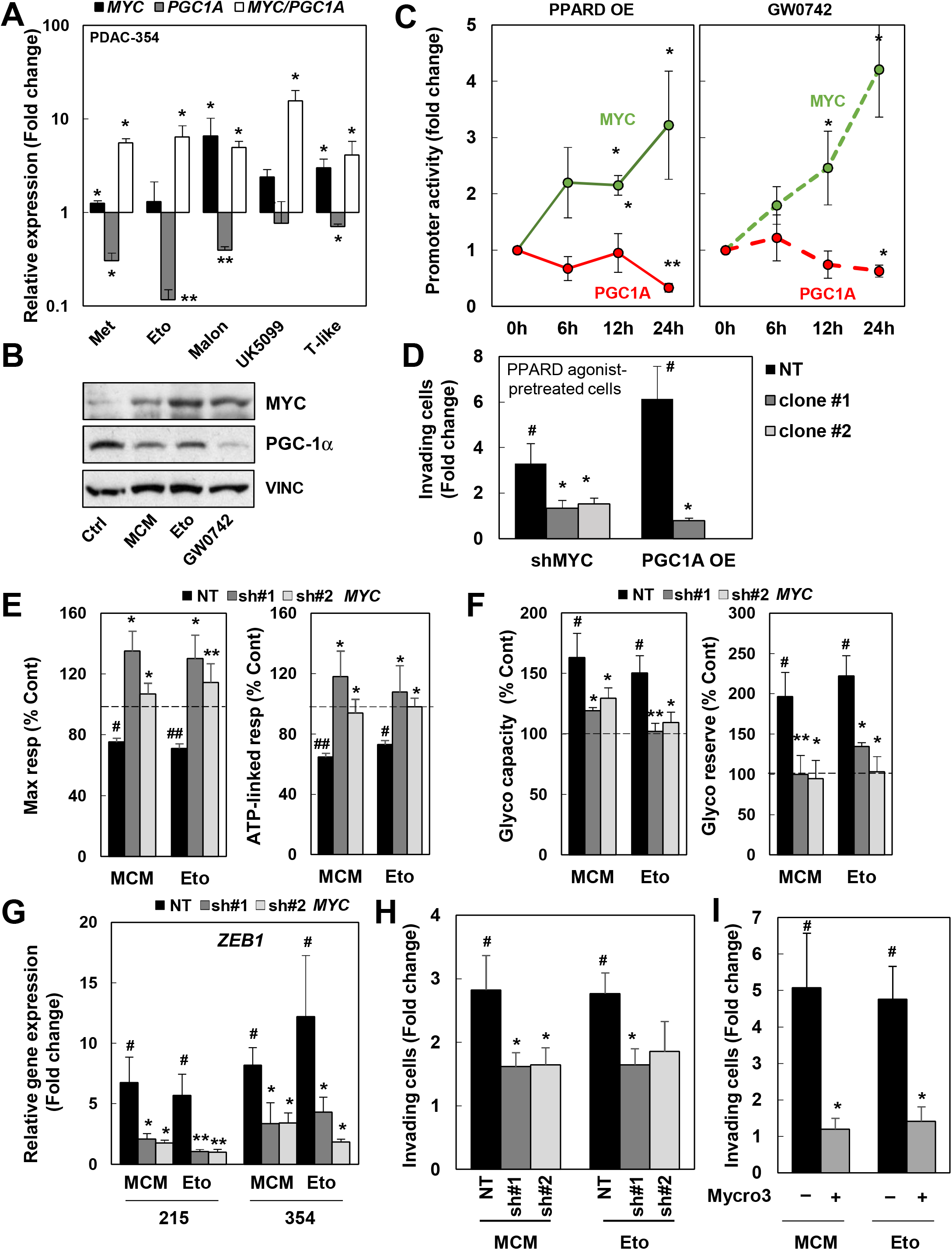
PPAR-δ rewires cellular metabolism regulating *MYC/PGC1A* balance. (**A**) Expression of *MYC, PGC1A* and *MYC/PGC1A* ratio in PDX-354 after mitochondrial energy deprivation during 48-72h. (**B**) MYC and PGC-1α expression measured by Western Blot following 48 h treatment with macrophage-conditioned medium (MCM), Etomoxir (Eto), or the PPAR-δ agonist GW0742 (5μM). Vinculin was used as loading control. (**C**) *MYC* and *PGC1A* promoter activity at the indicated times following treatment with PPAR-δ agonist GW0742 or *PPARD* overexpression (*PPARD* OE). (**D**) PDAC-354 cells were transduced with inducible lentiviral vectors expressing either a non-targeting shRNA (NT) or two different shRNAs against MYC (sh#1, sh#2) or the complete cDNA of *PGC1A*. Effect of MYC knockdown (shMYC, pooled data for sh#1 and sh#2) or PGC-1α overexpression (PGC1A OE) on invasiveness in response to treatment with 5μM PPAR-δ agonist L-165 for 48h. (**E-H**) PDAC-215 and 354 cells were transduced with inducible lentiviral vectors expressing either a non-targeting shRNA (NT) or two different shRNAs against MYC (sh#1, sh#2). Transduced cells were pre-treated with doxycycline for 48h and then incubated with MCM or Eto. (**E)** OCR changes for maximal respiration (left) and ATP-linked respiration (right). (**F)** Glycolytic capacity (left) and reserve (right). (**G)** *ZEB1* gene expression. (**H)** Invasive capacity. (**I**) PDAC-354 cells were treated with MCM or 20µM Eto for 48h in the presence or absence of the MYC/Max interaction inhibitor Mycro3 (25μM). Cells were then seeded in modified Boyden invasion chambers containing 20% FBS in the lower compartment and the number of invasive cells was assessed after 16h. All data are represented as mean ± SEM. # p<0.05, ## p<0.01, ### p<0.001 vs unstimulated control. * p<0.05, ** p<0.01, *** p<0.001 versus NT. See also Figure S.

To further corroborate our finding that the *MYC/PGC1A* ratio is crucially implicated in the metastatic process, we next analyzed different models with functional and/or physiological relevance. Specifically, we found that 1) the *MYC/PGC1A* ratio most closely correlated with patients’ survival (**Figure S6D**), 2) migrating cells showed both increased levels of EMT-associated genes and *MYC/PGC1A* ratio compared to non-migrating cells (**Figure S6E**); 3) the *MYC/PGC1A* ratio of disseminated cells from patients with advanced PDAC was 1,000–8,000x higher compared to the ratio observed for cells derived from primary tumors (**Figure S6F**), and 4) circulating tumor cells (CTC) in xenograft models showed increased *MYC* expression compared to the corresponding primary tumors (**Figure S6G**).

The latter finding was most pronounced in cells with increased expression of stemness genes as a putative pool of circulating CSCs (**Figure S6H**). Notably, *PGC1A* expression was very low to undetectable in most circulating single cancer cells and therefore the *MYC/PGC1A* ratio could not be calculated in this instance (data not shown). Interestingly, once the pro-metastatic PDX cells had formed actual liver metastases, they showed very low *MYC/PGC1A* ratios, with reduced *MYC* expression and *PGC1A* levels exceeding those found in primary tumors (**Figure S6I**). These changes in gene expression were associated with a reversion to their original mitochondria-driven metabolic state (**Figure S6J**).

To further validate the crucial functional role of the altered *MYC/PGC1A* ratio for inducing the EMT program, we next used an inducible *MYC* knockdown system to prevent *MYC* upregulation upon EMT induction. *MYC* knockdown was induced 48 hours before exposing the cells to EMT-inducing conditions. As expected, *MYC* knockdown essentially prevented the downregulation of *PGC1A* upon treatment with MCM or Etomoxir (**Figure S6K)** and the subsequent switch in the metabolic phenotype associated with EMT (**Figure 7E, 7F**). *MYC* knockdown also prevented *ZEB1* upregulation and induction of invasiveness (**Figure 7G, 7H**).

As blocking the *MYC/PGC1A*-governed metabolic program prevented the pro-metastatic phenotype induced by microenvironmental cues or fuel deprivation, we next aimed to pharmacologically inhibit *MYC* expression using the MYC/MAX interaction inhibitor Mycro3 in order to mimic the effects of *MYC* knockdown. Pretreating the cells with Mycro3 efficiently reduced the upregulation of *VIM* and *ZEB1* in response to EMT induction by MCM and Etomoxir (**Figure S6L**) and prevented induction of invasiveness (**Figure 7I**). These data could be further corroborated by overexpression of *PGC1A* prior to EMT induction, which prevented the metabolic changes induced by MCM and Etomoxir, respectively (**Figure S6M, left panel**) and consequently the cells did not acquire an invasive phenotype (**Figure S6M, right panel**).

Together, these data support our hypothesis that, upon PPAR-δ activation, *MYC* (through inhibition of *PGC1A*) not only governs the metabolic changes related to EMT, but also initiates and mediates the EMT/invasive program as a whole.

### Therapeutic targeting of PPAR-δ abrogates metastatic activity

Finally, we tested if PPAR-δ could be blocked pharmacologically to inhibit invasion and metastasis *in vitro* and *in vivo*. Indeed, pre-treatment with the PPAR-δ antagonists GSK0660 and GSK3787 or the inverse agonist DG172 inhibited the invasive capacity conferred by MCM or Etomoxir treatment (**Figure 8A**), or the basal invasive capacity of highly metastatic PDAC-265 cells (**Figure 8B**). Importantly, these *in vitro* results could be corroborated *in vivo* using a model of spontaneous metastasis following orthotopic injection of PDAC-265 cells. *PPARD* expression was significantly increased in PPAR-δ agonist GW0742-treated mice (**Figure 8C**), which translated into higher metastatic spread in GW0742-treated mice, whereas the PPAR-δ antagonist GSK3787 significantly reduced metastatic dissemination (**Figure 8D, 8E**). Of note, MYC and Vimentin protein expression were significantly increased in tumors treated with GW0742 (**Figure 8E, lower panel**).

**Figure 8.**
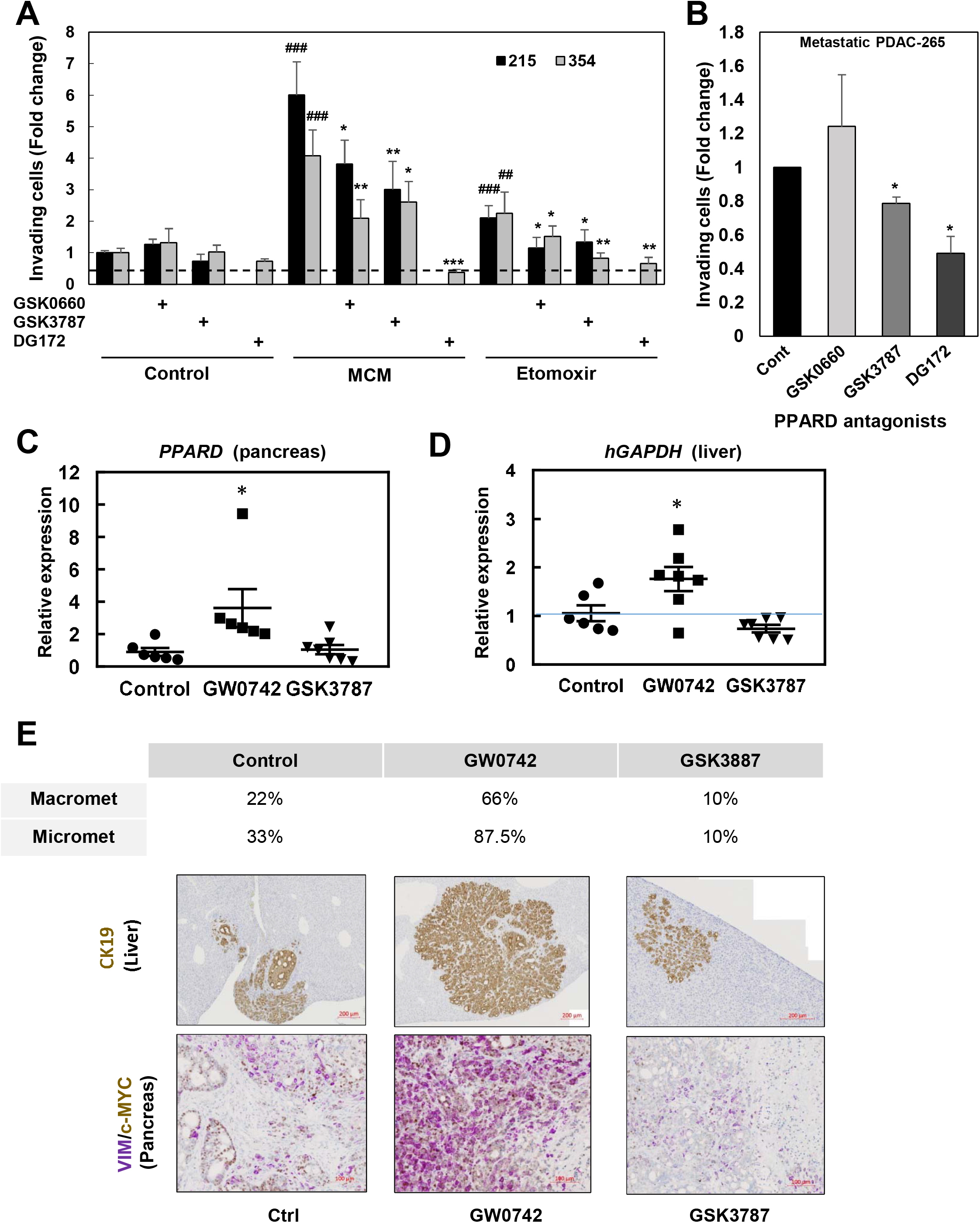
Therapeutic targeting of PPAR-δ impairs invasion *in vitro* and metastasis *in vivo*. (**A**) PDAC-215 and 354 cells were pre-treated with PPAR-δ antagonists GSK0660 (10µM) and GSK3787 (10µM) and inverse agonist DG172 (2.5µM) for 1h and then treated with MCM or Etomoxir for 48h. Invasion over 16h was assessed in modified Boyden invasion chambers. (**B**) Highly metastatic PDAC-265 cells were incubated with the PPAR-δ antagonists GSK0660 (10μM) and GSK3787 (10μM) and the PPAR-δ inverse agonist DG172 (1μM) for 48h and invasion was assessed after additional 16 hours. (**C, D**) Spontaneous metastasis upon orthotopic injection of 10^5^ metastatic PDAC-265-GFP-luc cells. Following implantation, mice were treated daily with either vehicle, the PPAR-δ agonist GW0724 (0.3mg/kg i.p.) or the PPAR-δ antagonist GSK3887 (3mg/kg i.p.) until termination of the experiment at week 9, when mice showed signs of disease. Tumor and onset of metastasis were assessed by weekly IVIS. (**C**) Expression of *PPARD* in pancreatic tumors measured by rt-QPCR. (**D**) Metastasis onset evaluated as *hGAPDH* absolute copy number. (**E**) Percentage of macro and micrometastases in the liver (**upper panel**). Expression levels of CK-19 in liver sections (**middle panel**, representative images), or c-MYC (brown) and VIM (purple) in pancreatic tumors was measured by IHQ (**bottom panel**, representative images). All data are represented as mean ± SEM. * p<0.05, ** p<0.01, *** p<0.001 vs NT cells; # p<0.05, ## p<0.01, ### p<0.001 vs control or single treatment.

In summary, PPAR-δ integrates nutrient-sensing and stromal signals to reprogram PDAC cell metabolism via *MYC/PGC1A*, promoting cancer cell invasiveness and *in vivo* metastasis in PDAC. Importantly, this process can be pharmacologically reversed using existing small molecule inhibitors, thus providing a potential new avenue for the treatment of advanced PDAC.

## Discussion

PPAR-δ is a member of the nuclear receptor superfamily of ligand-activated transcription factors. It regulates a variety of biological functions, in a cell and context-dependent manner, including cellular metabolism, proliferation, differentiation and survival, as well as inflammation (Giordano Attianese and Desvergne, 2015). Probably due to strong cell context-dependency and utilization of diverse model systems, the role of PPAR-δ in cancer has remained controversial (Wagner and Wagner, 2020). Although occasionally related to tumor suppression (Martín-Martín et al., 2018), increased *PPARD* expression has mostly been linked to enhanced metastasis in several *in vivo* models (Zuo et al., 2017). Even more importantly, poor patient outcome, including reduced metastasis-free survival correlated with *PPARD* expression in various cancer types (Abdollahi et al., 2007; Zuo et al., 2017). However, while accumulating evidence suggest that PPAR-δ also promotes tumor progression and metastasis in PDAC (Liu et al., 2020; Sanford-Crane et al., 2020; Zuo et al., 2017), other reports have questioned these finding (Coleman et al., 2013; Smith et al., 2016).

Our data now clearly support the notion that PPARD promotes progression and metastasis in PDAC. First, using single-cell analysis of various PDAC primary cultures, we found that *PPARD* was exclusively upregulated in cells actually undergoing EMT (**Figure 2A&B**). We also found PPARD to be overexpressed in the PAAD TCGA dataset (**Figure 3A**) and correlated with disease-free survival (**Figure 3B**). Interestingly, *PPARD*-high patients also showed enrichment for pathways related to cellular metabolism, inflammation, cell cycle, and EMT (**Figure 3D, 3E**), in line with our *in vitro* findings using single-cell analysis (**Figure 2 & S2**). Such broad transcriptional program controlled by PPAR-δ suggested to us that it represents a strong candidate for integrating the multiple pathways regulating tumor progression and metastasis in PDAC. Indeed, our present data now suggest that the implication of PPAR-δ in the interplay between tumor cells and TAMs may be bidirectional, since we found *PPARD* upregulation and activation in response to microenvironmental signals from TAMs (**Figure 2, 4A, 4B**). This could create a positive feedback loop *in vivo*, further promoting tumor progression via induction of EMT in cancer cells.

EMT can also be induced by metabolic stress resulting from impaired mitochondrial metabolism (**Figure 1**), which could be related to either genomic or transcriptional defects, e.g. lack of mitochondrial DNA (Guha et al., 2014), mutations in TCA cycle enzymes (Grassian et al., 2012; Loriot et al., 2012; Sciacovelli et al., 2016), or downregulation of components of the OXPHOS system (Gaude and Frezza, 2016). Our data now demonstrate that not only (epi-) genetic inhibition of mitochondrial function, but also functional inhibition of mitochondria, e.g. via pharmacological inhibition of the ETC, is sufficient to induce a metastatic program in PDAC, in line with findings for other cancer types (Han et al., 2018; Porporato et al., 2014).

Even more strikingly, we found that nutrient stress was similarly effective at inducing EMT and invasiveness of cancer cells (**Figure 1, S1**). Indeed, although glutamine deprivation has recently been described as an inducer of EMT via *Slug* upregulation in KPC-derived murine PDAC cells (Recouvreux et al., 2020), we here describe a wider phenomenon: inhibition of mitochondrial uptake of diverse carbon substrates (glutamine, pyruvate, fatty acids) and/or lack of oxygen, thereby mimicking the hypoxic and acidic tumor microenvironment, consistently induced EMT. Notably, mitochondrial activity was also repressed by microenvironmental signals such as TAM-derived factors, indicating a common metabolic route in the context of EMT (**Figure 5B, 5C**).

As PPAR-δ was the only PPAR family member activated by stromal signals or in response to metabolic stress conditions (**Figure 4B**), we hypothesized that PPAR-δ acts as an integrating sensor of diverse signals from the tumor microenvironment, and subsequently activates the EMT program to 1) gain metabolic plasticity and thereby adapt and survive in challenging environmental conditions and 2) acquire mobility, evade the primary tumor and search for more permissive environments elsewhere. Interestingly, it was recently proposed that inflammatory signals can trigger a pseudo-starvation response driving invasiveness, independent of nutrient abundance (García-Jiménez and Goding, 2019), suggesting that PPAR-δ controls such starvation/pseudo-starvation responses as a prerequisite for induction of EMT and subsequent metastasis in PDAC.

To our knowledge, this is the first report associating PPAR-δ with tumor progression and metastasis via metabolic rewiring. PPAR-δ initiates a global pro-metastatic metabolic program via increasing the *MYC/PGC1A* ratio. Notably, changes in the ratio predicted the aggressiveness of PDAC cells and overall patient survival more accurately than each of the two genes individually. Indeed, the pro-invasive effects of enhanced PPAR-δ activation could be reversed by either *MYC* knockdown/pharmacological inhibition or *PGC1A* overexpression (**Figures 7, S7**). Pharmacological or genetic induction of *PPARD* resulted in a rapid upregulation of *MYC* (24h), suggesting a direct interaction as the *MYC* promoter carries a PPAR responsive element (PPRE; Genecard), although *MYC* upregulation may occur indirectly via the microRNA Let-7c (Shah et al., 2007). But our promoter activity assays confirmed that *PPARD* stimulation directly induced *MYC* promoter activity and subsequently reduced *PGC1A* promoter activity (**Figure 7C**). The resulting pro-invasive effects could be reversed by knockdown of *MYC* or overexpression of *PGC1A* (**Figure 7D**).

MYC and PGC-1α have been connected to metabolic switch and tumor progression/metastasis. Specifically, *MYC* expression promotes cellular de-differentiation, EMT, and increased metastatic potential (Bian et al., 2017; Ischenko et al., 2015; Soucek et al., 2013). Indeed, the molecular signature of aggressive squamous/mesenchymal PDAC includes *MYC*-activated signaling pathways (Bailey et al., 2016). Moreover, analysis of PDAC models revealed that *MYC* overexpression is associated with less differentiated tumors and a glycolysis-related gene signature (Bian et al., 2017). Indeed, we found that *MYC* upregulation suppressed *PGC1A* resulting in an altered metabolism with enhanced global glycolytic/plastic capacity (conferred by *MYC*), accompanied by inhibition of mitochondrial oxygen consumption and activity (as a result of *PGC1A* downregulation via *MYC*).

Our data show that metabolic plasticity is crucial to support the increased energetic needs during invasion and the subsequent metastatic process. As metastatic cells decrease their mitochondrial function (Danhier et al., 2013; Schafer et al., 2009), they need to rely on alternative sources to maintain their energy balance, and activation of glycolysis seems to be the most plausible option. However, our results demonstrate that the increase in glycolytic activity upon EMT induction is rather modest (**Figure 5B, 5C and S5**). Instead, we found that glycolytic reserve was more profoundly enhanced in EMT cells, rather suggesting increased metabolic plasticity and diversification of metabolic substrates, e.g. alternative sugars or fatty acids (**Figure 5A, S6**).

Although CSCs and non-CSCs are similarly capable of undergoing EMT, regardless of their basal metabolic phenotype (**Figure S1E**), CSCs are the most capable and aggressive cells for establishing new metastatic sites due to their inherent self-renewal and tumor-initiating capacities (Hermann et al., 2007). Previously we also reported that most CSCs in the primary tumor are strictly dependent on OXPHOS activity and that these harbor the highest tumorigenic potential. Here we now expand on these findings by showing that in CSCs undergoing EMT the self-renewal capacity remained essentially unchanged (**Figure S1F**). While these results were rather unexpected they do suggest an intricate interplay between stemness, EMT and cellular metabolism (Daniel et al., 2021). Considering the importance of maintaining stemness in cancer (Wang et al., 2019; Zhang et al., 2017), we hypothesize that during the EMT process PPAR-δ becomes a key driver of stemness, rendering CSCs less dependent on mitochondrial metabolism. Future studies should further dissect this potential mechanistic duality in CSCs.

Finally, we found that genetic or pharmacological targeting of PPAR-δ inhibited tumor aggressiveness and metastasis *in vitro* and *in vivo* (**Figure 8**). These data are in line with previous reports for murine PDAC models showing that *Ppard* knock-down strongly decreased tumorigenesis mouse melanoma cells (Zuo et al., 2017), and *Ppard* knockout inhibited tumor progression in KC mice on a high-fat diet (Liu et al., 2020). Taken together, accumulating data now strongly support the concept that PPAR-δ inhibition reduces the *MYC/PGC1A* ratio and thereby diminishes PDAC progression and metastasis. These data provide the rational for developing novel PPAR-δ-targeting treatment strategies to combat advanced pancreatic cancer.

## Supporting information

Supplemental Experimental Procedures

## Author Contributions

B.P-A, D.B. and S.T. acquired, analyzed, interpreted data and assisted in the development of the study concept; S.C., S.C.-S., Q.Z., J. T., M.C., Z.G., P.I., L.P.-P., S.M.C., S.L.-E. and P.E. acquired and analyzed *in vitro* data; A.C-G, Q.Z., J. T., M.C., Z.G., performed single-cell RNA extraction and ddPCR and 10x RNA-seq, respectively; L.R.-C., S.C., P.J., and B.S. Jr performed *in vivo* experiments; U.L. and M-L.L. performed bioinformatics analyses; M.Y. analyzed GC-MS data; A.L. contributed with funds and critical advice; C.H. and P.S. developed the study concept, obtained funding, interpreted the data, and wrote the manuscript.

## Acknowledgements

Authors would like to acknowledge the use of the BCI Flow Cytometry and Pathology Facilities, as well as the CIBA Flow Cytometry, Pathology and Microscopy Facilities (Servicios Científico-Técnicos, IACS-Universidad de Zaragoza). We would like to thank Arkaitz Carracedo, Veronica Torrano, and Natalia Martin-Martin for constructive data discussion and providing us with the *PGC1A* and *PPARD* overexpression lentiviral vectors.

The research was supported by the ERC Advanced Investigator Grant (Pa-CSC 233460, to C.H.), the European Community’s Seventh Framework Programme (FP7/2007-2013) under grant agreement n° 602783 (CAM-PaC, to C.H.), the Fondazione del Piemonte per l’Oncologia – IRCCS (PTCRC-Intra 2021, to C.H.), the Pancreatic Cancer Research Fund (2015 Award round, to P.S.), the Instituto de Salud Carlos III through the Miguel Servet Program (CP16/00121 to P.S.) and Fondo de Investigaciones Sanitarias (PI17/00082 to P.S.) (both co-financed by European funds (FSE: “el FSE invierte en tu futuro” and FEDER: “una manera de hacer Europa,” respectively), the Worldwide Cancer Research Charity together with Asociación Española contra el Cáncer (AECC) (19-0250, to P.S.), the Fero Foundation (to B.S.,Jr.), a Coordinated grant (GC16173694BARB) from the AECC (to B.S., Jr.), an Investigator Grant (NHMRC #1196405, to U.S.) and a Cancer Council NSW project grant (RG20-12, to U.S).

## Experimental Procedures

### Primary human PDAC cells

For primary cultures, PDAC tissue fragments were minced, enzymatically digested with collagenase (Stem Cell Technologies) for 90 min at 37°C (Mueller et al., 2009), and after centrifugation for 5 min at 1,200 rpm the pellets were resuspended and cultured in RPMI, 10% FBS, and 50 units/ml penicillin/streptomycin. For experiments, cells were cultured in DMEM:F12 supplemented with B-27, L-Glutamine (all from Gibco, Life Technologies), 50 U/mL penicillin–streptomycin (Sigma) and β-FGF (PeproTech). PDXs tissues were obtained through the Biobank of the Spanish National Cancer Research Centre (CNIO), Madrid, Spain (references M-20/002-1, I409181220BSMH, 1204090835CHMH) and the ARC-NET Biobank at the ‘Rossi’ University of Verona Hospital, Italy (reference 6.B.04 - Samples PDAC-10953). Cancer cells from advanced PDAC patients were isolated and expanded from peripheral blood (Shanghai Jiaotong University School of Medicine, Protocol No 20130905), as previously described (Agerbæk et al., 2018).

### Primary human macrophages and conditioned media

Leucocyte cones from anonymous healthy donors were obtained from the National Blood Transfusion Service (UK) according to City and East London Research Ethics Committee (17/EE/0182). Cones were stored at 4°C and used within 24 hours of delivery to maintain cell viability. Monocyte-derived human macrophage culture, polarization into M2-like macrophages and generation of conditioned medium were as previously described (Sainz et al., 2014, 2015). Monocyte-derived human macrophage cultures were maintained in IMDM (Gibco) supplemented with 10% human AB serum and polarized by incubation with 0.5ng/ml of macrophage colony-stimulating factor for 48 hours (MCSF; PeproTech). To generate conditioned media, macrophages were then washed with PBS and cultured for additional 48 hours in supplemented DMEM:F12 (see previous section). Media was then collected, centrifuged and supernatant stored at -80°C.

### Single-cell capture, library Preparation, and RNA-seq

The samples (ETO-treatment vs CTRL, MCM-treatment vs CTRL) were labeled with Cell Hashing antibodies following the manufacturer’s instruction (BioLegend), cells were counted on Countess II automated cell counter (Thermo Fisher) after staining, and up to 25,000 cells were loaded per lane on 10X Chromium microfluidic chips (10X Genomics). Single-cell capture, barcoding, and library preparation were performed using the 10X Chromium Single Cell 3’ Reagent Kits version 3 chemistry, and according to the manufacturer’s protocol (#CG000185). cDNA and HTO libraries were checked for quality on the Agilent 4200 Tapestation, and quantified by KAPA qPCR before sequencing on a single lane of a NovaSeq 6000 S4 flow cell (Illumina) to an average depth of 100,000 reads per cell.

### Single-Cell Data Processing, Quality Control, and Analysis

The Cell Ranger pipeline (v1.3, 10X Genomics) was used to firstly convert Illumina base call files to FASTQ files, then demultiplexing was conducted before aligning FASTQs to the GRCh38 genome reference and producing the digital gene-cell counts matrix. Samples were combined using the Cell Ranger aggregate function, which merges output from multiple runs to normalized to the same sequencing depth before generating a gene-barcode (cell) expression matrix. Potential doublets were identified by DoubletFinder (McGinnis et al., 2019) and removed before proceeding to downstream analysis. Quality control, normalization, clustering, dimensionality reduction and visualization were performed using R toolkit Seurat package (Butler et al., 2018). Gene-cell matrices were filtered to remove cells with fewer than 500 unique molecular identifiers (UMI) counts and 500 detected genes, and cells with more than 15% mitochondrial gene counts were also filtered. The gene set enrichment analysis was conducted using ssgsea function from GSVA package. RNA-seq data are available at NCBI dbGaP under the accession number GSE184871.

### XF extracellular flux analysis

Single-cell suspensions from trypsinized secondary spheres/adherent cultures were plated in XF96 Cell Culture Microplates previously coated with Cell-Tak (BD Biosciences) at a cellular density of 30,000 cells/well. For OCR determination, cells were incubated in base assay medium supplemented with 2mM glutamine, 10mM glucose, and 1mM pyruvate for 1h, prior to the measurements using the XF Cell Mito Stress Kit. Concentrations of oligomycin and FCCP were adjusted for each primary cell type. For glycolytic metabolism measurements, cells were incubated in basal media supplemented with 2mM glutamine and 1mM pyruvate prior to injections using the Glycolysis Stress Test kit. Experiments were run in a XF96^e^ analyzer, and raw data were normalized to protein content. Unless indicated otherwise, all reagents and materials were from Agilent Seahorse XF Technologies (Agilent Technologies).

### Invasion assay

Invasion assays were performed using 24-well 8.0μm PET membrane invasion chambers coated with growth factor reduced Matrigel™ (Corning). After 48h of pre-treatment, 10^5^ primary PDAC cells were seeded to coated inserts in serum free media. Invasion towards 20% FBS was tested after 12-24h incubation at 37°C in a humidified atmosphere of 5% CO_2_. Invaded cells were fixed with 4% paraformaldehyde, stained with DAPI and imaged on the Olympus Fluorescence microscope (model BX51). Cell number was analyzed using automated ImageJ particle analysis software.

### *In vivo* metastasis and treatments

For classical metastasis assay upon intrasplenic injection, pre-treated PDAC-354 CMV-Luciferase-RFP-TK expressing cells were re-suspended in 50μl of Matrigel and injected in the spleen of NSG mice (NOD Scid interleukin (IL)-2 receptor γ chain knockout mice; Charles Rivers) at a concentration of 0.5×10^5^ cells per injection. After 7 days, splenectomy was performed. For spontaneous metastasis assay, PDAC-265 cells were re-suspended in 30μl of Matrigel and injected orthotopically to NSG mice at a concentration of 1×10^5^ cells per injection. Mice were then imaged weekly using the IVIS Spectrum Imaging System (Caliper Life Sciences). Mice were anaesthetized with isoflurane (2%) and injected intraperitoneally with 150 mg/kg of luciferin (Caliper Life Sciences) diluted at 15 mg/mL in PBS. For the experiment shown in Figure 5D, mice were treated for three consecutive days with GW0742 (0.3 mg/kg i.v.) after surgery. For the experiment shown in Figure 5F, mice were treated with oral doxycycline (2mg/ml drinking water) and Etomoxir (15 mg/kg, i.p. every day) for 7 days after intrasplenic implantation. For the experiment shown in Figure 8E, mice were treated daily with either vehicle (PBS), the PPAR-δ agonist GW0724 (0.3 mg/kg i.p.) or the PPAR-δ antagonist GSK8337 (3 mg/kg i.p.) until termination of the experiment. Once a minimum of 1×10^6^ ROI bioluminescence in liver was achieved in at least 3 mice after 5 minutes following injection, or if signs of ascites developed, all experimental mice were sacrificed (9 weeks). Livers and pancreas were harvested, imaged on collection and fixed in 4% PFA. Procedures were conducted in accordance with institutional and national regulations (Animals in Science Regulation Unit, Home Office Science, London, UK; Project License PPL70/8129; Ethical Conduct in the Care and Use of Animals as stated in The International Guiding Principles for Biomedical Research involving Animals (Council for International Organizations of Medical Sciences (CIOMS)); Universidad de Zaragoza Ethics Committee; project licenses PI22/17 and PI41/20).

### Statistical analysis

Results for continuous variables are presented as means ± SEM unless stated otherwise. Treatment groups were compared with the independent samples t test. Pair-wise multiple comparisons were performed with the one-way ANOVA (two-sided) with Bonferroni adjustment. p values < 0.05 were considered statistically significant. All analyses were performed using Prism GraphPad (version 5.04).

**Further description of experimental procedures is provided as supplemental information**.

