## Supplemental Figures for "PPAR-delta acts as a metabolic master checkpoint for metastasis in pancreatic cancer"

### Figure S1, related to Figure 1

**Mitochondrial energy deprivation promotes epithelial-to-mesenchymal transition in PDAC cells. (A) Left panel**: representative micrographs of parental PDAC-215 and their metformin-resistant counterparts. **Right panel**: *Vimentin* (*VIM*) and *ZEB1* expression in Metformin-resistant cells (215M, 253M, 354M) *vs* treatment naïve parental cells (PDAC-215, 253, 354). (**B**) PDAC-215, 253, and 354 cells were treated for 48h with the indicated concentrations of Metformin, Malonate, UK5099, or hypoxia (3% $O_2$) with or without glutamine deprivation (low Q, 0.2 mM) and low pH (50 µM HCl). Expression of EMT-associated genes was determined by rtPCR. Data were normalized to HPRT. (**C**) Cells were exposed for 72h to normoxia (20% $O_2$) or hypoxia (3% $O_2$) ± the following metabolic stress conditions : low glutamine (low Q, 0.2mM), low glucose (low Glc, 1mM), low pH (50µM HCl) or combinations thereof (low Q & low pH; low Glc & low pH). Pooled data for PDAC-185, A6L, 215, 253, and 354 (n≤4 for each cell type). (**D**) *VIM* and *ZEB1* expression in cells co-cultured for 72 h with M2-differentiated human primary macrophages (M0) or human primary pancreatic stellate cells (PSC). Pooled data from PDAC-253 and 354 (n=2 for each cell type). (**E**) *ZEB1* expression for cells sorted for CD133 and mitochondrial content (MitoTracker) and treated for 48h as indicated. Pooled data from PDAC-253 and 354. (**F**) Sphere formation after 7 days of treatment with Control, MCM, and Etomoxir. Some of the data shown here are also included in the pooled data graphs represented in Figure 1. Data are represented as mean ± SEM. * p<0.05, ** p<0.01. *** p<0.001. & p<0.05 *vs* M0

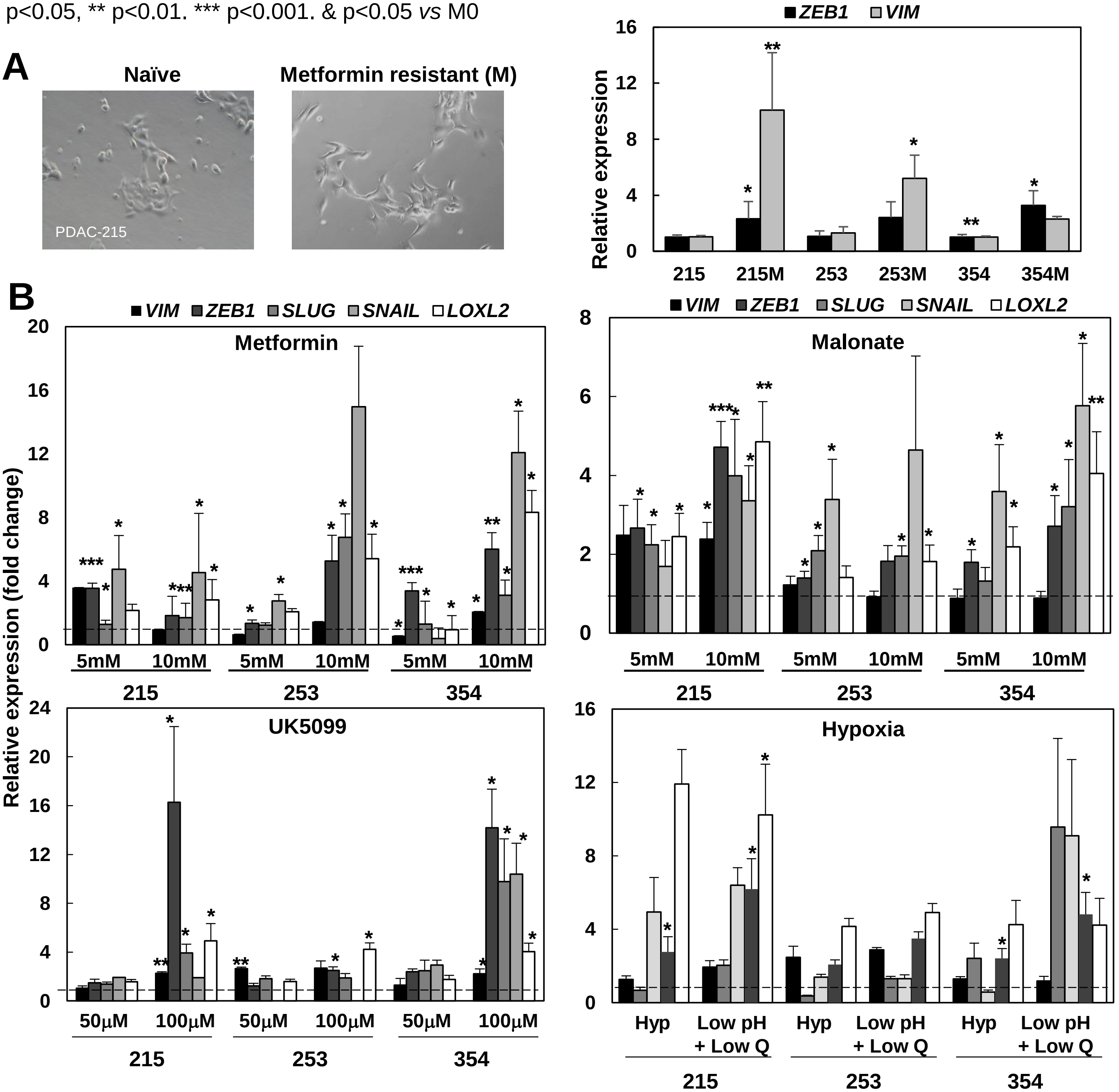

### Figure S1, related to Figure 1

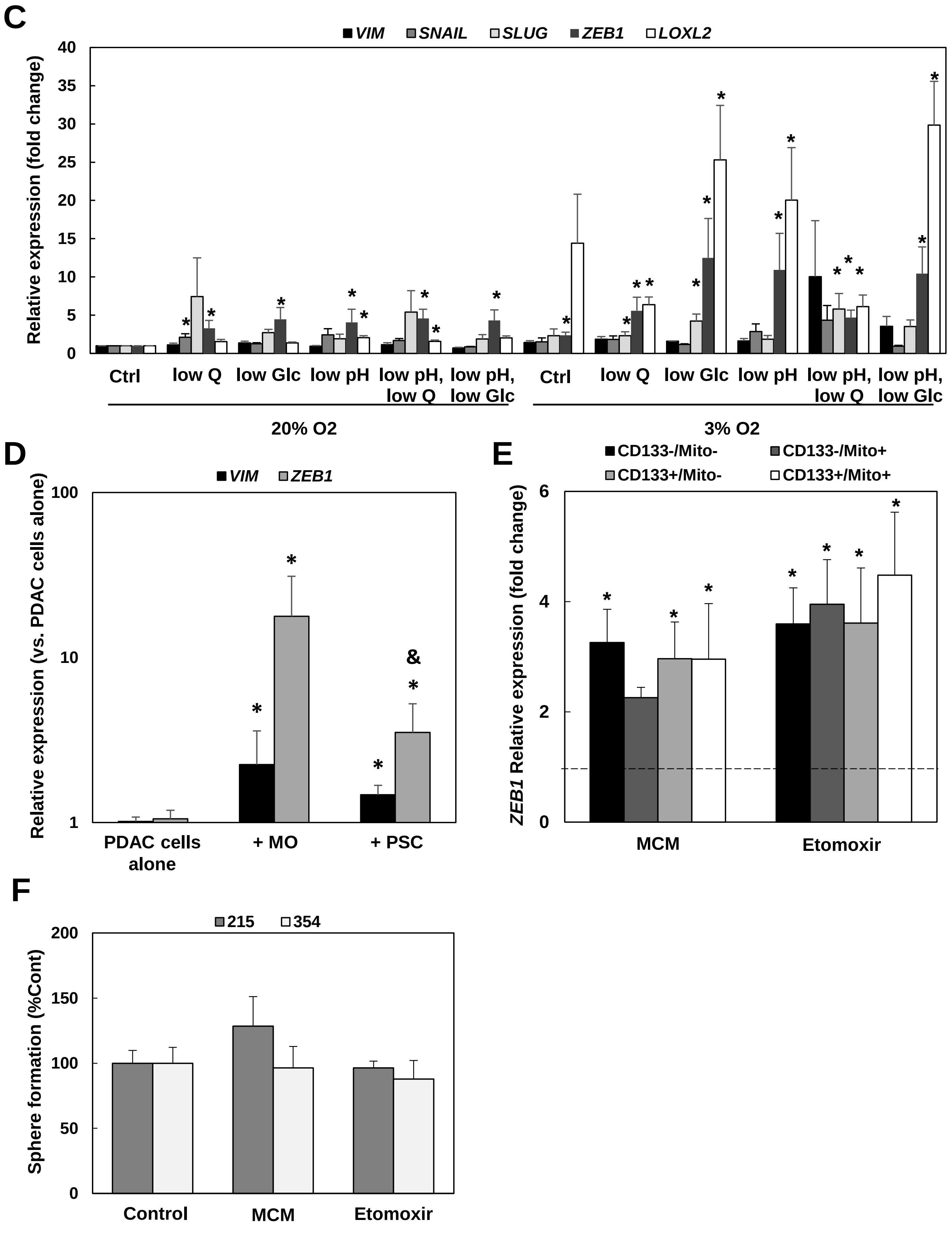

### Figure S2, related to Figure 2

**Transcriptomic analyses reveals a common transcriptional program associated to EMT**. Cells were treated with control vehicle (CTRL), macrophage-conditioned medium (MCM), or 20µM Etomoxir (ETO) for 48h. (**A**) representative images. (**B**) Unsupervised clustering of viable PDAC cells exposed to CTRL, MCM or ETO, subjected to single-cell RNAseq (10X Chromium platform) and represented as UMAP plots. (**C**) Gene set enrichment analysis (GSEA, Hallmark gene set) for the Hallmark Hypoxia in the different transcriptional clusters identified by single-cell RNAseq. (**D**) Heat map showing scaled normalized expression of differentially expressed genes between control and MCM cells, with cells as columns and genes as rows. Genes were selected based on absolute log2FoldChange > 0.8 and p<0.05. (**E**) Heatmap of top differentially expressed genes (absolute log2FoldChange > 1.5, padj < 0.1) after analysis by bulk RNAseq. Hierarchical clustering of genes differentially expressed in control cells versus treated cells. (**F**) Commonly up-regulated and metabolism-related pathways as determined by GSEA (Hallmark gene set) for Etomoxir-treated PDAC cells.

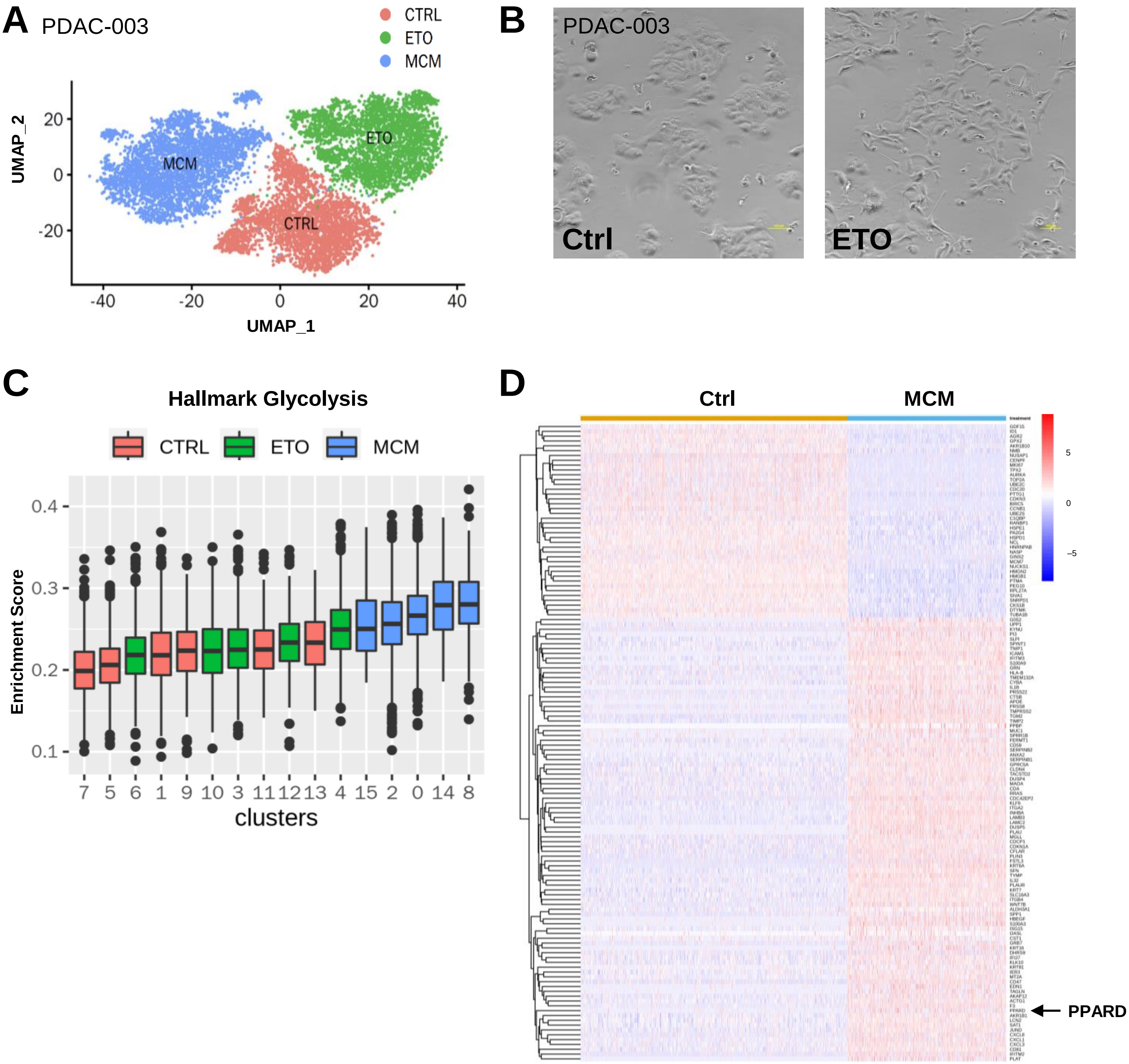

### Figure S2, related to Figure 2

E

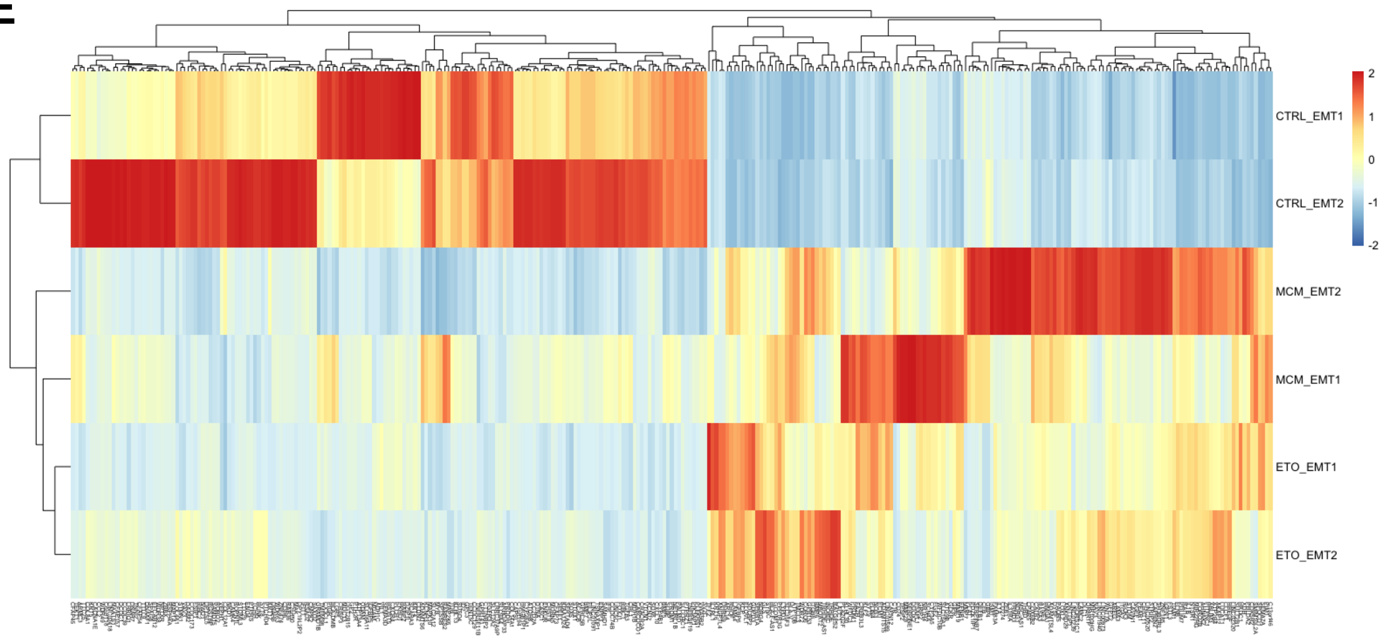


F

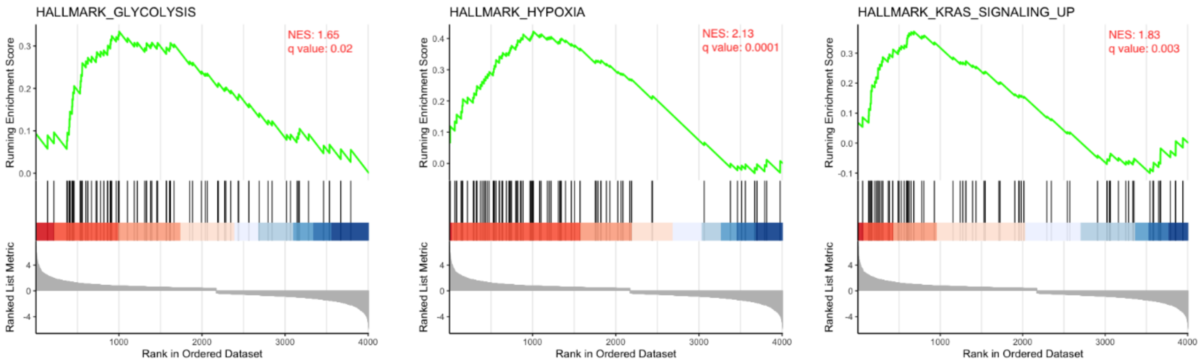

### Figure S3, related to Figure 4

**Activation of PPARD induces EMT and metastasis.** (**A**) Gene (24 & 48h; left panel) and protein (48h; right panel) expression for the PPAR family members following treatment with macrophage-conditioned medium (MCM), Etomoxir (Eto), or the PPAR-δ agonist GW0742 (5 μM). (**B**) Representative gene expression kinetics of the indicated genes after treatment with MCM (left) of Etomoxir (right) in PDAC-215 cells. (**C**) Induction of PPAR targets after 24h of treatment with MCM, Etomoxir or the PPARD agonist GW0742 (10μM). (**D**) Preferential induction of PPARD after 72h of treatment with the PPARD agonists GW0742, GW501516, and L-165 (each 10μM). Pooled data from PDAC-A6L, 185, 215, 253, and 354 cells.

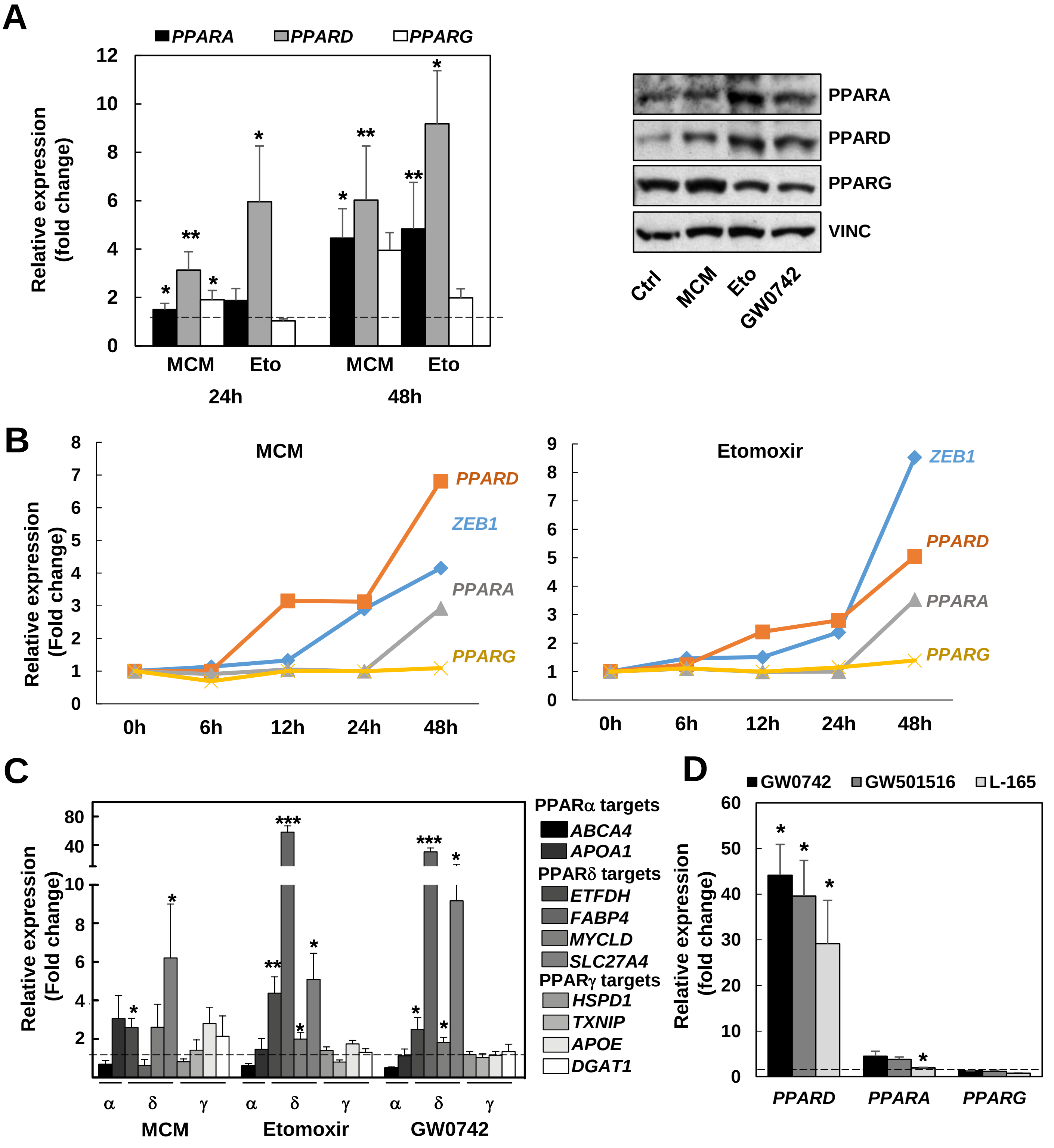

### Figure S4, related to Figure 4

**PPAR-δ controls the EMT program induced by microenvironmental and nutrient-sensing signals**. (**A**) Changes in *PPARD* and EMT-associated genes after 72h of treatment with the indicated concentrations of the PPAR-δ agonists GW0742, GW501516, and L-165. Pooled data from PDAC-A6L, 185, 215, 253, and 354 cells. (**B**) Lack of changes in gene expression (left panel, pooled data for PDAC-215, 253, and 354) and morphology (right panel, PDAC-354 cells) treated with WY14643 (PPAR-α agonist), Rosiglitazone (PPAR-γ agonist), and GW0742 (PPAR-δ agonist) (each 10μM). (**C, D**) PDAC-253, 215, and 354 cells transduced with inducible lentiviral vectors expressing either a non-targeting shRNA (NT) or three different shRNA against PPARD (sh#1, sh#2, sh#3). Cells were pre-treated with doxycycline for 24 hours to induce shRNA expression and subsequently incubated with macrophage-conditioned medium (MCM), Etomoxir (Eto) or the PPAR-δ agonist L-165. Changes in expression of *PPARD* (**C**) and EMT-related genes (**D**) for PDAC-253 (upper panel), 215 (middle panel) and 354 (lower panel) cells. Data are represented as mean ± SEM. # p<0.05, ## p<0.01, ### p<0.001 vs respective NT control.

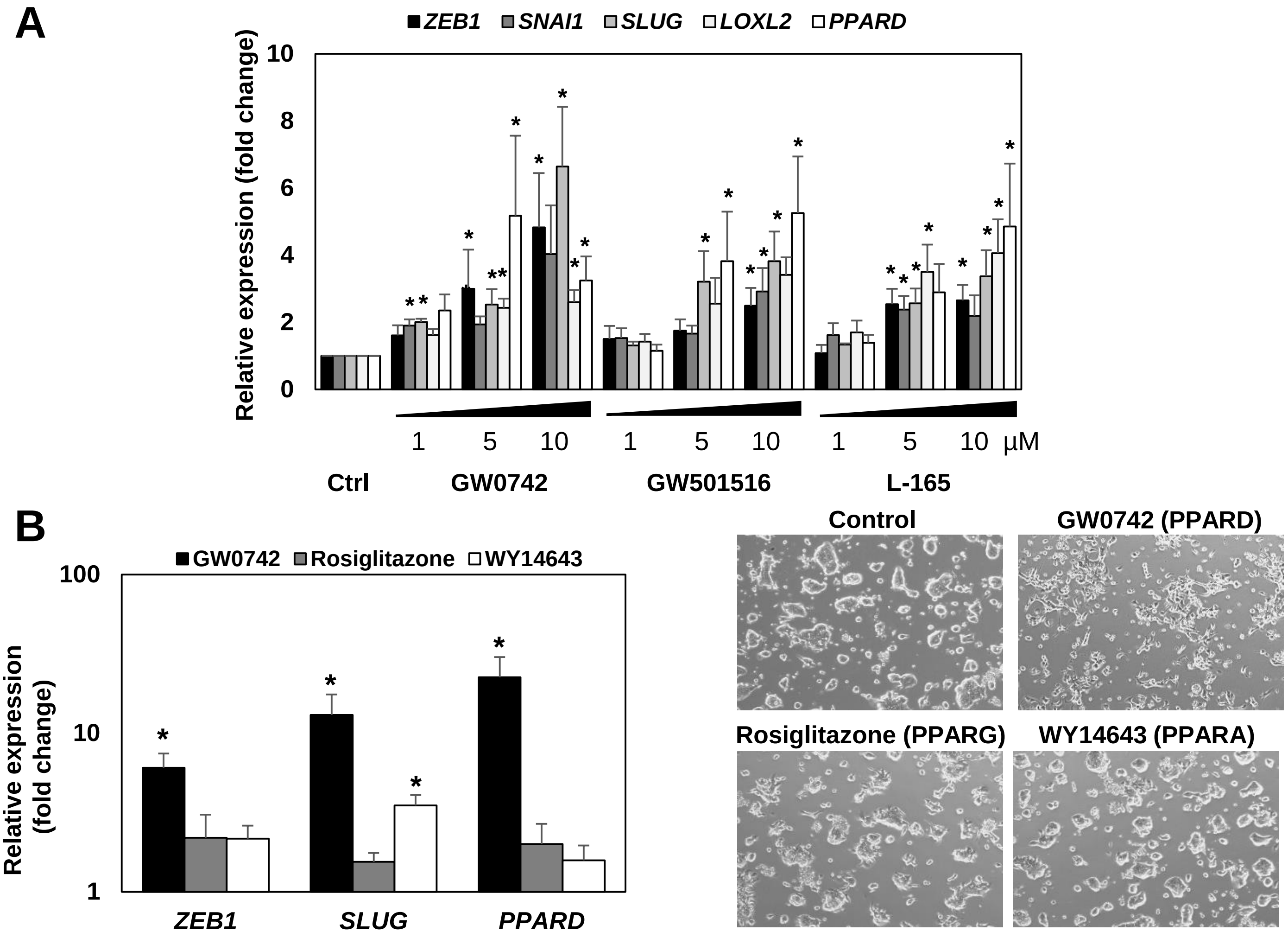

### Figure S4, related to Figure 4

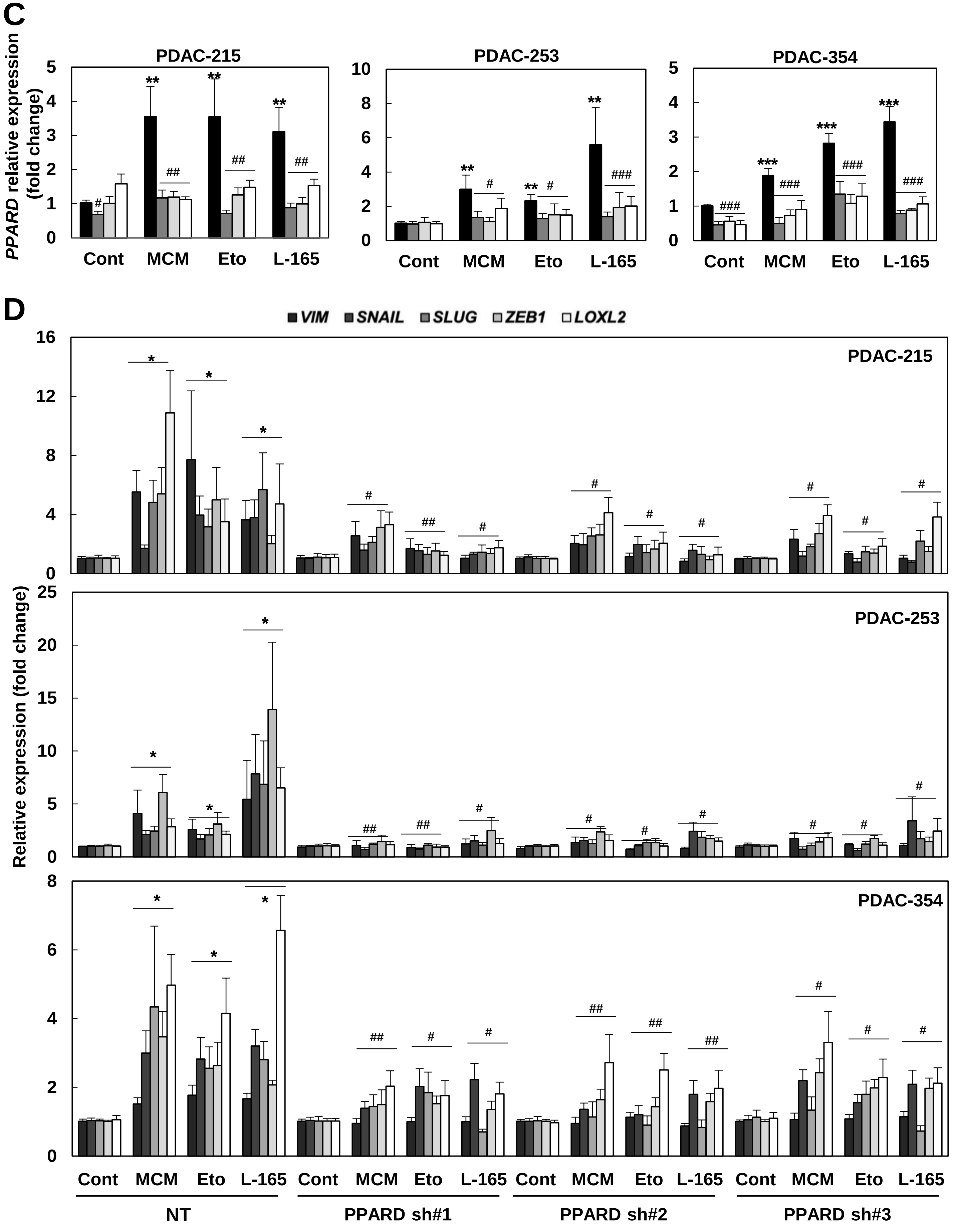

### Figure S5, related to Figure 5

**EMT induced by microenvironmental and nutrient-sensing signals relates to common metabolic changes.** **(A)** OCR and (**B**) ATP-linked OCR for PDAC spheres cultured alone (alone), together with M2 macrophages (+MO) or with PSC (+PSC). Pooled data for sphere-derived PDAC-253 and 354 cells. (**C**) Maximal and ATP-linked respiration in non-CSC or CSC alone or co-cultured with MO. Pooled data from PDAC-253 and 354 cells. (**D**) Cells incubated for 48h with macrophage-conditioned medium (MCM) or 20mM Etomoxir (Eto). Representative FACS for NBDG uptake in PDAC-253 cells, (**E**) Lactate concentration in supernatants, (**F**) $^{13}$C-Lactate and $^{13}$C-Alanine in supernatants as measured by GCMS after incubation with $^{13}$C-Glucose for 12h. All data are represented as mean ± SEM. * p<0.05, ** p<0.01, *** p<0.001.

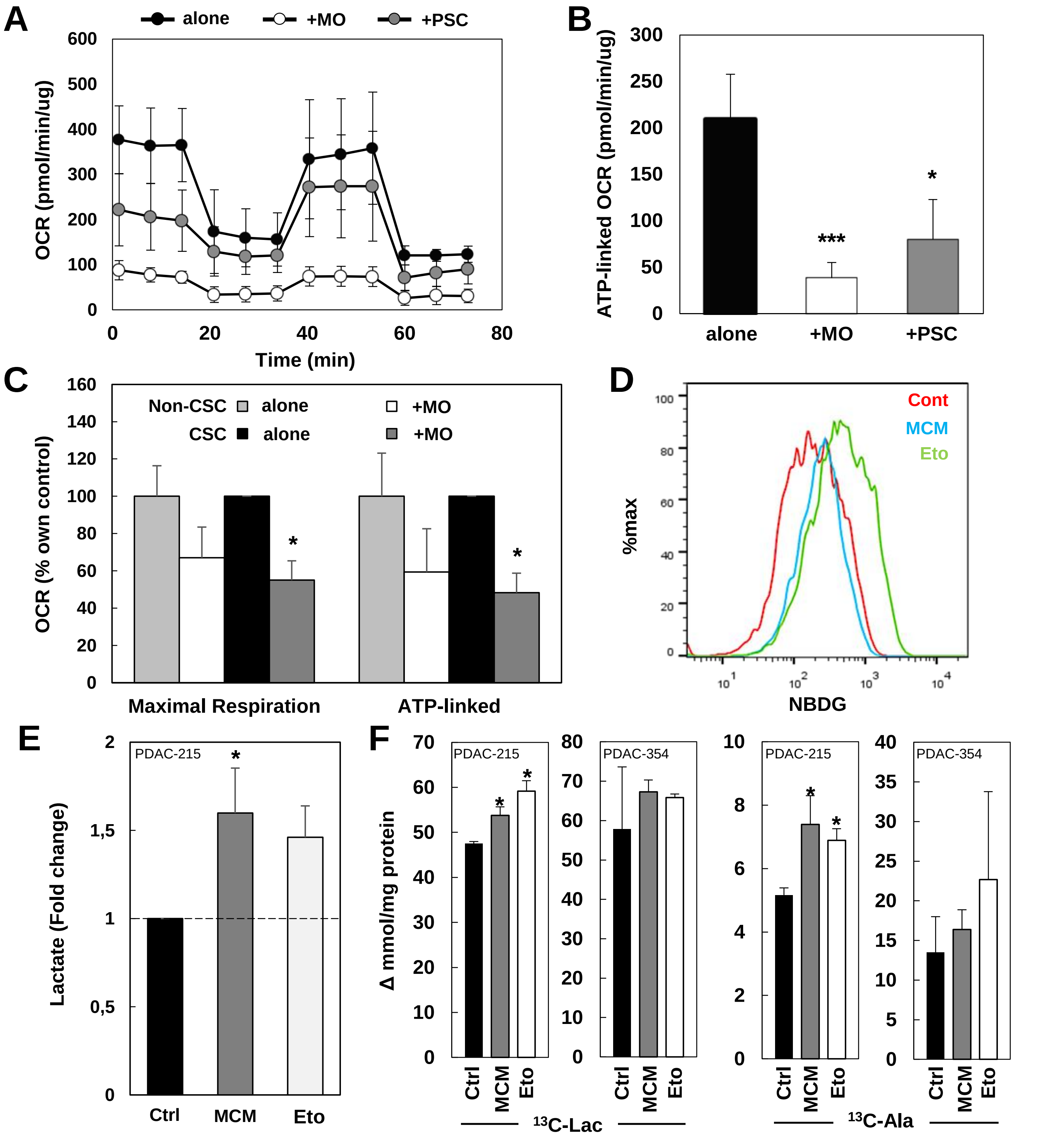

### Figure S6, related to Figure 7

***PPARD* controls cellular metabolism via *MYC/PGC1A* ratio to promote EMT and metastasis**. Where indicated, cells were treated with macrophage-conditioned medium (MCM) or Etomoxir 20 μM (Eto) for 48h. Cells transduced with lentiviral constructs were pretreated with doxycycline for 48h before treatments. (**A**) MYC overexpression promotes EMT-like phenotype**.** Left panel: representative images of parental and MYC-overexpressing PDAC-215 cells (MYC OE). Right panel: *MYC*, *VIM*, and *ZEB1* expression. (**B**) Expression of *MYC*, *PGC1A* and *MYC/PGC1A* ratio in PDAC-253 (**left**) and PDAC-215 (**right**) after mitochondrial energy deprivation during 48-72h. (**C**) PDAC-253, 215, and 354 cells transduced with a non-targeting shRNA (NT) or three different shRNA against PPARD (sh#1, sh#2, sh#3) were incubated with MCM or Eto or the PPARD agonist L-165 to determine *MYC/PGC1A* expression. (**D**) Patients from the TCGA dataset were dichotomized for the tumor expression levels for *MYC*, *PGC1A* and *MYC/PGC1A* ratio. Kaplan Meier survival curves for disease-free survival are shown. (**E**) EMT-related gene expression of cells that had transmigrated towards 20% FBS for 16h versus gene expression of non-transmigrated cells. Pooled data from PDAC-253, 10953, and 354 (n=2 for each cell type). (**F**) *MYC*, *PGC1A* and *MYC/PGC1A* ratio comparing six local disease PDX (A6L, 185, 215, 253, 354, 10953) vs 5 metastatic PDX (SiC-002, 004, 006, 021, and 023). (**G, H**) Single GFP+ PDAC-354 cells sorted from the blood (CTC) and pancreas (primary) of mice bearing orthotopic tumors were analyzed for the expression of *MYC* and *PGC1A* (**G**) or stratified into CSC versus non-CSC based on their expression of pluripotency-related genes (*NANOG*, *KLF4*, *SOX2*, *OCT4*) (**H**). (**I**) *MYC*, *PGC1A* and *MYC/PGC1A* in primary tumors and liver metastasis from mice bearing orthotopic tumors. (**J**) *In vivo* metabolomics data after $^{13}$C-Glucose injection comparing primary tumors with liver metastases. (**K**) PDAC-215 and 354 cells transduced a non-targeting shRNA (NT) or two different shRNAs against MYC (sh#1, sh#2) were incubated with MCM or Eto to measure *MYC* (**left**) and *PGC1A* (**right**) expression. (**L**) *VIM* and *ZEB1* expression in PDAC-354 cells treated with MCM (**left**) or Eto (**right**) in the presence or absence of the MYC/MAX interaction inhibitor Mycro3 (25μM). (**M**) PDAC-354 cells transduced with an inducible construct for PGC-1α overexpression were incubated with MCM or Eto to test maximal respiration (**left**) or invasion (**right**). All data are represented as mean ± SEM. * p<0.05, ** p<0.01, *** p<0.001 vs NT cells; # p<0.05, ## p<0.01, ### p<0.001 vs control or single treatment.

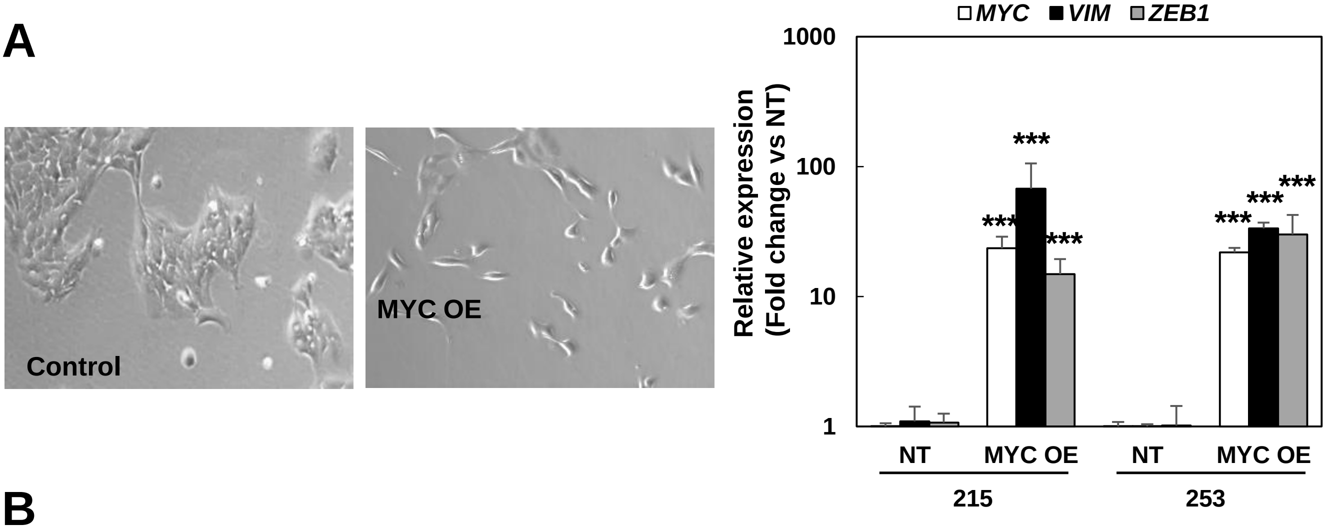


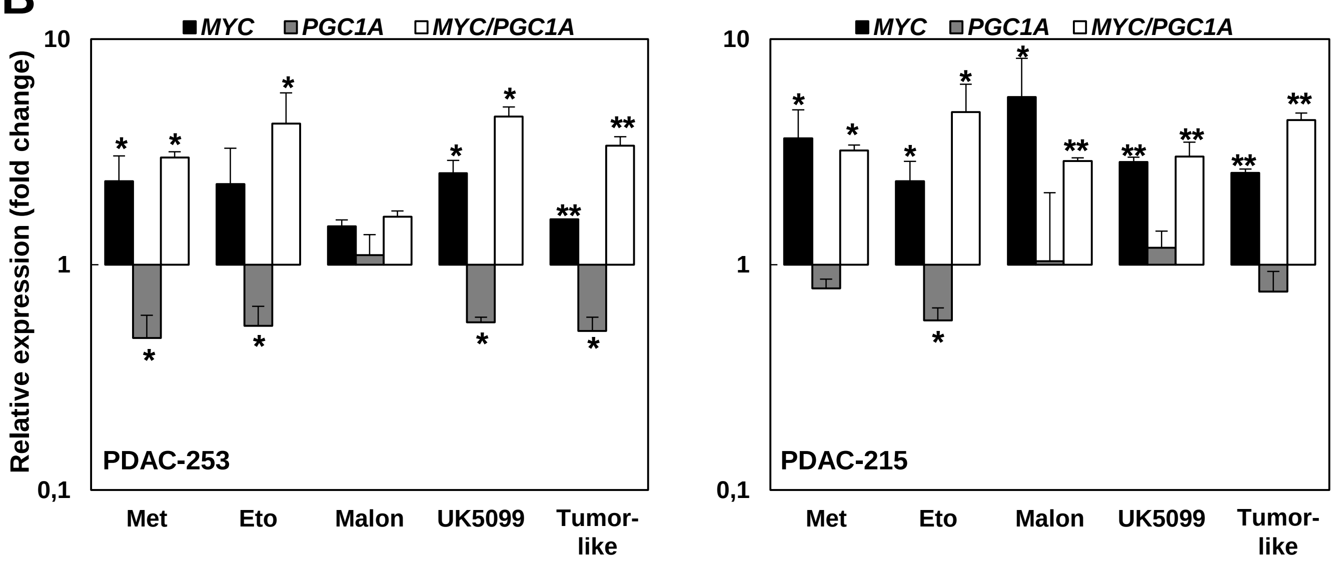

### Figure S6, related to Figure 7

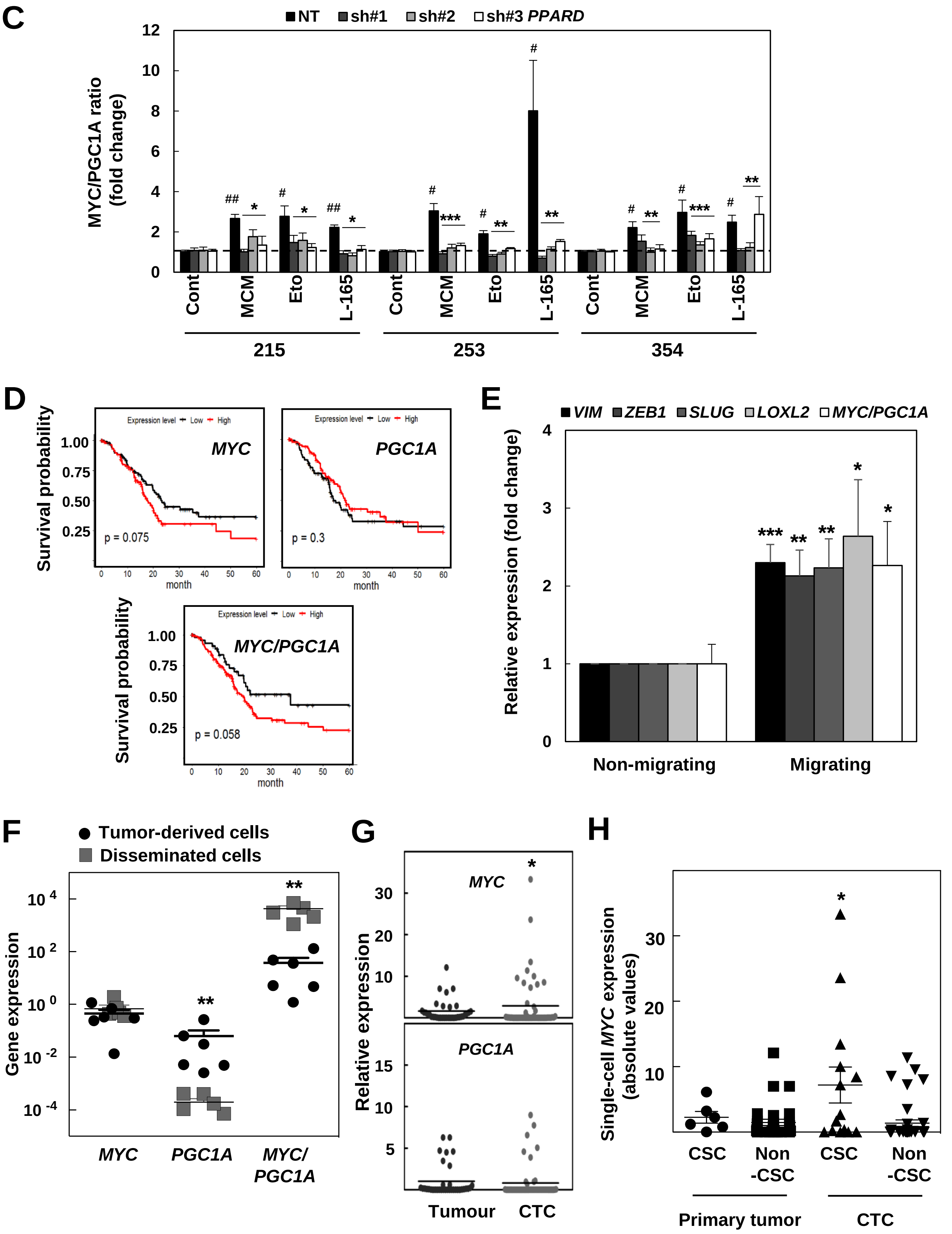

### Figure S6, related to Figure 7

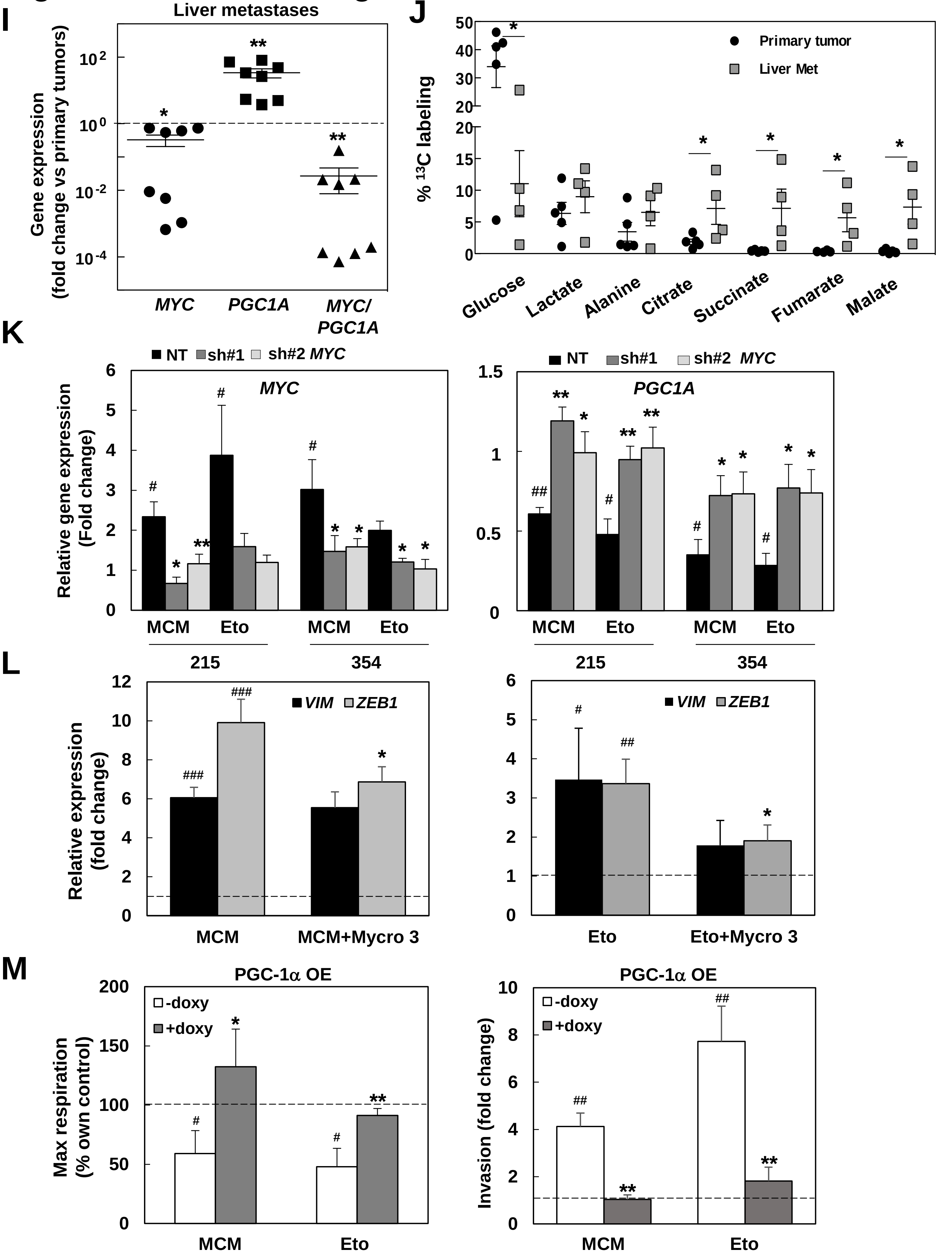
