## Supplemental Experimental Procedures for "PPAR-delta acts as a metabolic master checkpoint for metastasis in pancreatic cancer"

**Chemicals.** The mitochondrial inhibitors Metformin, Malonate and Etomoxir (all from Sigma) were dissolved in water and used at the different doses indicated in figure legends. The MPC inhibitor UK5099 (Merck chemicals) was dissolved in DMSO and used at 50 and 100 μM. Doxycycline (Sigma) dissolved in water was used at different doses to induce the expression of cDNA or shRNA. PPAR-δ agonists GW501516 (Generon), GW0742 (Cayman) and L165,041 (Cayman) were dissolved in DMSO and used at 1, 5 and 10 μM. PPARA and PPARG agonists (WY14643, Rosiglitazone respectively; both from Cayman) were dissolved in DMSO and used at 10 μM. PPAR-δ antagonists GSK0660 (Sigma) and GSK3787 (Cayman) were dissolved in DMSO and used at 10 μM. The PPAR-δ inverse agonist DG172 (Tocris Bioscience) was dissolved in DMSO and used at 1 μM. The Myc-Max heterodimerization/DNA binding inhibitor Mycro3 (Aobious) was dissolved in DMSO and used at 25 μM

**Flow Cytometry and Cell Sorting.** Primary pancreatic cells were resuspended in Sorting buffer (1× PBS; 3% FBS [v/v]; 3μM EDTA [v/v]) before analysis with a LSR Fortessa Cell Analyser platform (BD Bioscience) or sorting with a FACSARIA Fusion Cell Sorter (BD Bioscience). To identify pancreatic CSCs anti-CD133/1-PE (Miltenyi) or appropriate isotype-matched control antibody were used. CSCs were further subdivided depending on their mitochondrial content after staining (10 min, 37°C) with 100nM Mitotracker Deep Red (Life Technologies). Alternatively, cells were incubated for 20 minutes at 37°C with 100mM 2-NBDG (2-(N-(7-Nitrobenz-2-oxa-1,3-diazol-4-yl)Amino)-2-Deoxyglucose (Life Technologies) prior to FACS analysis to evaluate glucose uptake. DAPI was used for exclusion of dead cells. Data were analyzed with FlowJo 9.2 software (Tree Star Inc., Ashland, OR).

**Co-culture of PDAC and macrophages or CAFs.** $10^5$ M2-like polarized macrophages or primary CAFs were seeded to pre-soaked 6 well 0.4μm permeable polycarbonate membrane transwell (Corning) in IMDM supplemented with 10% Human Serum (Sigma). In parallel, 1.5x $10^5$ primary PDAC cells were seeded in 6 well adherent plates (adherent cultures) or ultralow attachment plates (sphere cultures) in supplemented DMEM:F12. Following attachment overnight, macrophages transwells were added to wells containing PDAC cells, and co-cultures were maintained in supplemented DMEM:F12 for 4 days.

**Cancer stem cell-enriching culture.** PDAC spheres were generated and expanded in supplemented DMEM-F12 (see Primary Human PDAC cells section). A total of $10^4$ cells/ml was seeded in ultra-low attachment plates (Corning) as described previously (Gallmeier et al., 2011). For serial passaging, spheres were harvested at day 7 using a 40μm cell strainer, dissociated to single cells with trypsin, and then re-grown in the same conditions for 7 days.

**PPAR activity assay.** PPAR specific DNA binding activity was performed on nuclear extracts upon 24 hours of treatment using the PPAR (alpha, delta, gamma) Transcription Factor Assay Kit (Abcam), following manufacturer´s instructions.

**Lactate production.** After treatments, cell culture supernatants were treated following manufacturer's instructions (Lactate Assay Kit II, Sigma).

**Lentiviral constructs.** For knockdown experiments, the shERWOOD UltramiR Lentiviral Inducible pZIP target gene set for MYC (TLHSU2300-4609) or PPARD (TLHSU2300-5467) were used (Transomic Technologies, Hunsville, AL). For overexpression, HA-PGC1A or HA-PPARD were cloned into a TRIPZ inducible plasmid as previously described (Martín-Martín et al., 2018; Torrano et al., 2016). Lentiviruses were generated by co-transfection of 293T cells with respective lentivirus shuttle backbone vectors, the Pax2 packing plasmid and the vesicular stomatitis virus glycoprotein expression plasmid (pCMV-VSV-G) and pTAT using Lipofectamine 2000 (Invitrogen) according to the manufacturers' protocol. Supernatants were collected 48h post-transfection, filtered through a 0.45μm-pore-size filter (BD Biosciences), aliquoted, frozen, and subsequently tittered using 293T cells and flow cytometry.

**MYC and PGC1A promoter reporter assays.** HEK293 cells were plated in 6-well plates at a density of 250.000 cells / well. Cells were transfected 24 hours later with 1 μg of plasmid PGC-1A promoter reporter vector (Active motif S722424), HBM-luc (gift from Linda Penn (Facchini et al., 1997), Plasmid #35155 (Addgene) or control promoter vector (Active motif 32021) and, when indicated, 1 μg of PPARD-overexpression vector using lipofectamine 2000 (Invitrogen) according to the manufacturers' protocol. After 8 hours, PPAR-δ was induced by treatment with 5 μM GW0742 or overexpression with 1 ng/ml doxycycline, and luciferase expression was read 48 hours later using LightSwitch Assay Reagent (Active motif LS010) according to manufacturer's protocol.

**Western blot.** After treatment, cells were lysed in RIPA buffer (R0278, Sigma-Aldrich, St. Louis, MO, USA) supplemented with protease inhibitors (J64156) and phosphatase inhibitors (J61022) (Alfa Aesar, Thermo Fisher Scientific, Haverhill, USA). Proteins were quantified using the Pierce™ BCA Protein Assay Kit (Thermo Fisher Scientific). Extracts were submitted to electrophoresis in 10% Tris-Glycine gels (XP001002, Invitrogen), transferred to a PVDF membrane (88518, Thermo Fisher Scientific) and incubated overnight at +4°C with the different primary antibodies (Table S1). After washes with PBS-Tween 0.1%, the membranes were incubated with peroxidase-conjugated goat anti-rabbit or goat anti-mouse secondary antibodies. Bound antibody complexes were detected using Pierce™ ECL Western Blotting Substrate (32109, Thermo Fisher Scientific) and visualized on CL-X Posure™ Films (34091, Thermo Scientific).

| Protein | Company | Reference | Clone | Dilution |
|---|---|---|---|---|
| **PPAR-α** | Santa Cruz Biotechnology | sc-398394 | H-2 | 1:1000 |
| **PPAR-δ** | Santa Cruz Biotechnology | sc-74517 | F-10 | 1:1000 |
| **PPAR-γ** | Santa Cruz Biotechnology | sc-7273 | E-8 | 1:1000 |
| **c-MYC** | Cell Signaling Technology | 9402S | | 1:1000 |
| **PGC-1α** | Santa Cruz Biotechnology | sc-13067 | H300 | 1:1000 |
| **Vinculin** | Sigma Aldrich | SAB4200080 | V-284 | 1:2000 |
| **β-Actin** | Sigma Aldrich | A2228 | AC-74 | 1:10000 |

*Table S1. List of primary antibodies used for Western blot*

**Immunohistochemistry.** Formalin-fixed paraffin-embedded (FFPE) mice livers and pancreata were serially sectioned (3μm thick). CK19, c-MYC or vimentin antibodies (Table

S2) were applied for 1 hour at room temperature. After incubation, slides were washed in PBS and incubated with horseradish peroxidase-conjugated avidin (ABC Standard: Vector Laboratories). Antigen was visualized using 3,3'diaminobenzidine (DAB+) chromogen (DakoCytomation) for 2 minutes and then sections were counterstained in Mayer Haematoxylin for 2 minutes. Histological quantification of digitalized slides was performed using ZEN blue edition (Zeiss). Following manual optimization of antibody, automated staining was achieved using the Ventana Classic Automated system.

| Protein | Company | Reference | Clone |
|---|---|---|---|
| **c-MYC** | ABCAM | ab32072 | Y69 |
| **CK-19** | DAKO | IR615/IS615 | RCK108 |
| **Vimentin** | DAKO | IR630 | V9 |

*Table S2. List of primary antibodies used for IHC*

**RNA preparation and RTqPCR.** Total RNA was isolated using TRI Reagent following manufacturer's instructions. For tissue, a piece of approximately 5 mg tissue from a pancreatic tumor or a liver lobe was mechanically homogenized in TRI Reagent. One microgram of purified RNA was used for cDNA synthesis using the QuantiTect Reverse Transcription Kit (Qiagen) followed by SYBR green RTqPCR using an Applied Biosystems Viia-7 real-time thermocycler (Applied Biosystems). The list of utilized primers is shown in the table below. For Figure 3, a commercially available PCR array for 188 carbohydrate metabolism-related genes was used (Prime PCR Assay, 384 format, Bio-Rad, Hercules, CA). For this assay, PCR reactions were run with the SsoAdvanced universal SYBR Green Supermix (Bio-Rad).

| Gene | Forward Primer | Reverse Primer |
|---|---|---|
| ***HPRT*** | TGACCTTGATTTATTTTGCATACC | CGAGCAAGACGTTCAGTCCT |
| ***LOXL2*** | GGCACCGTGTGCGATGACGA | GCTGCAAGGGTCGCCTCGTT |
| ***C-MYC*** | CCCGCTTCTCTGAAAGGCTCTC | CTCTGCTGCTGCTGCTGGTAG |
| ***PGC1A*** | TGACTGGCGTCATTCAGGAG | CCAGAGCAGCACACTCGAT |
| ***PPARA*** | CTGGAAGCTTTGGCTTTACG | ACCAGCTTGAGTCGAATCGT |
| ***PPARD*** | CTCTATCGTCAACAAGGACG | GTCTTCTTGATCCGCTGCAT |

| | | |
|---|---|---|
| ***PPARG*** | GACCTGAAACTTCAAGAGTAC | TGAGGCTTATTGTAGAGCTGAGTC |
| ***SLUG*** | ATGCCGCGCTCCTTCCT | TGTGTCCAGTTCGCT |
| ***SNAIL*** | GCTCCTTCGTCCTTCTCCTC | TGACATCTGAGTGGGTCTGG |
| ***VIM*** | GACAATGCGTCTCTGGCACGTCTT | TCCTCCGCCTCCTGCAGGTTCTT |
| ***ZEB1*** | GTTGATGAATGCGAGTCAGATGC | CTGGTCCTCTTCAGGTGCC |
| ***GFP*** | GCAAAGACCCCAACGAGAAG | TCACGAACTCCAGCAGGACC |
| ***ABCA4*** | AGAATAACCGGACGCTGCTC | TCACCAAACCGGGCATAGAC |
| ***APOA1*** | TGCCCACTCTATTTGCCCAG | CTCACTGGTCCTGGCAATGT |
| ***ETFDH*** | GGACAGTCCTCCTGTTGTGT | TTCCATGTTCACTCCCAGGCA |
| ***FABP4*** | TGGGCCAGGAATTTGACGAA | CACATGTACCAGGACACCCC |
| ***MYLCD*** | GTCCGGGAAATGAATGGGGT | GCCAGGTAACCCGTTCTAGG |
| ***SLC27A4*** | GGCTCAGGGGCCAATAAACT | ACAGATGAGGCGGGTCAATG |
| ***HSPD1*** | CCTGCACTCTGTCCCTCACTC | GGTCTCATCTGGCGAAAGACT |
| ***TXNIP*** | CAACTTGCTGCCCGACAAAA | TGGGTGGCATGCAAGGTATT |
| ***APOE*** | GTTGCTGGTCACATTCCTGG | GCAGGTAATCCCAAAAGCGAC |
| ***DGAT1*** | TCGCCTGCAGGATTCTTTAT | GCATCACCACACACCAGTTC |
| ***hGAPDH*** | TCCTGTTCGACAGTCAGCCGC | ACGACCAAATCCGTTGACTCCG |
| ***mGAPDH*** | TGCACCACCAACTGCTTAGC | TCTTCTGGGTGGCAGTGATG |
| ***ZsGreen*** | | |

*Table S3. List of primers used for qPCR*

**Single-cell expression analysis by ddPCR from an orthotopic *in vivo* PDAC model.** In order to generate a PDAC mouse model with spontaneous metastasis, $5x10^4$ $GFP^+$ PDAC-354 cells were orthotopically injected into NU-Foxn1nu (Charles River) nude mice. CTC content was checked by FACS every two weeks from a small blood sample. Once mice showed high CTC count or signs of disease, they were humanely sacrificed and pancreas and total blood were harvested and processed for FACS sorting. For blood samples cell solution was resuspended in ACK lysing buffer (Thermo Fisher Scientific) during 5min, and then washed with PBS. Pancreatic tumors were minced mechanically with a scalpel and enzymatically digested with collagenase P for 15min at 37°C followed by trypsin for 3 min.

After pancreatic tumor digestion and blood processing, cell suspensions were blocked in Flebogamma (Grifols) for 15min at 4°C and incubated with anti-hEpCAM-APC antibody (Miltenyi Biotec) and an appropriate isotype-matched control antibody (IgG2a-APC, BD

Bioscience) for 30min at 4°C. Then, cells were washed in PBS and resuspended in sorting buffer [1X PBS; 3% FBS (v/v); 3mM EDTA (v/v); 2mg/ml DAPI]. Single cells double positive for hEpCAM-APC and GFP positives were FACS sorted (BD FACSAria™ II) in 96-well plates containing 100µl of GTC mix [4.19M Guanidine thiocyanate; 25mM Na Citrate pH=7.3; 15mM Sarcosyl; 11mM 2-Mercaptoetanol; 18µM Glycogen], and plates were stored at -80°C until RNA extraction was performed.

For single cell RNA extraction, each cell previously resuspended in 100 µl of GTC, was mixed with 10 µl of 2 M NaOAc pH=4 and 100 µl 100% Phenol/Chloroform pH=4. After centrifuging each sample at 13,000 rpm and at 4°C for 10 min, the upper aqueous phase was transferred to low binding tubes and mixed with 2 µl of 5mg/ml Linear Acrylamide (Amresco®). Finally, 400 µl of 100% Isopropanol were added to each sample, and samples were stored at -20°C overnight. The next day, samples were centrifuged at 13,000rpm and at 4°C for 30 minutes. After that, pellets were washed twice, first with 80% ethanol and then with 100% ethanol, by centrifuging at 13,000 and at 4°C for 10 minutes after each wash. Finally, pellets were air-dried and incubated with 7µl of RNase-free water for 1 hour to ensure the correct dissolution of RNA. Samples were stored at -80°C or immediately reverse-transcribed.

Next, reverse transcription reaction was directly run with the complete product of extraction measurement was used for cDNA synthesis using SuperScript® VILO cDNA Synthesis Kit (Thermo Fisher Scientific) following manufacturer's instructions. After that, a pre-amplification reaction with AmpliTaq Gold® 360 Master Mix (Thermo Fisher Scientific) was carried out with forward and reverse primers of every gene of interest (see below).

Finally, single-cell gene expression was performed using the QX100™ Droplet Digital™ PCR system (Bio-Rad) following the official ddPCR™ application guide. First, a mix containing the pre-amplified cDNA, a primer pair for one of the genes of interest, QX200™ ddPCR EvaGreen Assay and QX200™ Droplet Generation Oil for EvaGreen was used for droplet generation inside the QX100™ Droplet Generator (Bio-Rad). After droplet generation, 40µl of the generated droplet emulsions were transferred to a new 96-well PCR plate (Eppendorf), foil sealed (PX1™ PCR Plate Sealer, Bio-Rad) and amplified in a C1000 Touch™

Thermal Cycler (Bio-Rad) according to Bio-Rad recommendations. Finally, signal was quantified in the QX100™ Droplet Reader and analyzed using the software QuantaSoft 1.3.2.0. to calculate the concentration of the target DNA sequence (copies/μl), estimated by modelling as a Poisson distribution. Differences in gene expression were evaluated with Student's t-test or one-way ANOVA applying Bonferroni adjustment. $P<0.05$ was considered as statistically significant.

| Gene | Forward Primer | Reverse Primer |
|---|---|---|
| ***HPRT*** | TGACCTTGATTTATTTTGCATACC | CGAGCAAGACGTTCAGTCCT |
| ***C-MYC*** | CCCGCTTCTCTGAAAGGCTCTC | CTCTGCTGCTGCTGCTGGTAG |
| ***PGC1A*** | TGACTGGCGTCATTCAGGAG | CCAGAGCAGCACACTCGAT |
| ***NANOG*** | AGAACTCTCCAACATCCTGAACC | TGCCACCTCTTAGATTTCATTCTCT |
| ***OCT4*** | CTTGCTGCAGAAGTGGGTGGAG | CTGCAGTGTGGGTTTCGGGCA |
| ***KLF4*** | ACCCACACAGGTGAGAAACC | ATGTGTAAGGCGAGGTGGTC |
| ***SOX2*** | AGAACCCCAAGATGCACAAC | CGGGGCCGGTATTTATAATC |

*Table S4. List of primers used for ddPCR.*

**Bulk RNAseq analysis**

The paired-end RNA-seq libraries were generated using TruSeq Stranded mRNA kits with 200 ng of total RNA per sample. After quality control, reads were aligned to the human genome build 38 and GENCODE gene annotation human release 27 using STAR (Dobin et al., 2013) (version 2.5.3a). The number of reads that map to each gene were quantified by HTSeq (Anders et al., 2015), (Version 0.9.1). Differential expression analysis was carried out using R/Bioconductor package DESeq (Love et al., 2014) (version 3.4.4). The RNA seq data was deposited at Gene Expression Omnibus (GSE135686). Differentially expressed genes identified in the macrophage and Etomoxir treatment were subjected to unsupervised hierarchical clustering using Pearson correlation distance matrix and complete linkage.

Gene set enrichment was performed using GSEA software (Subramanian et al., 2005; Mootha et al., 2003) from broad institute using the Hallmark gene set database (Liberzon et al., 2015).

**Human data analysis**

Expression data from human PDAC tissue and normal tissue were analyzed using the webserver GEPIA2 (TCGA and the GTEx project databases; http://gepia2.cancer-pku.cn/) (Tang et al. 2019). The Pearson correlation coefficient was calculated to study the association of *PPARD* gene with the EMT-related genes *ZEB1*, *SNAIL* and *SLUG*. For disease-free survival analysis, the Hazard Ratio (HR) was calculated using the Cox Proportional Hazards model for PDAC from the respective upper and lower quartiles of expression of *PPARA*, *PPARD* and *PPARG*. The samples included in the top and bottom quartile of *PPARD* expression in the TCGA dataset were compared in GSEA. The GSEA module of the Genepattern suite from the Broad Institute was used, with 1000 permutations and FDR < 25% was considered statistically significant.

**GC-MS Metabolomics**

For *in vitro* metabolomic tracing experiments, second-generation sphere-derived cells and adherent cells were seeded in sphere medium. 3 hours before starting the metabolic labeling, cells were equilibrated in unlabeled tracing medium (DMEM without pyruvate, with 10mM glucose, 2mM L-glutamine, Pen/Strep, bFGF and B-27). At time 0, the medium was substituted with 4ml/sample of tracing medium with 10mM U-$^{13}$C6-Glucose (Cambridge Isotope Laboratories). Medium samples were collected for analysis at the indicated time points. After 24h the cells were washed with cold PBS and scrapped in dry ice-cold MeOH for metabolite extraction. Cells were lysed in 3:1 methanol:chloroform by pulse sonication for 1 hour. Water was added to make an extraction solvent ratio of 3:3:1 methanol:chloroform:water. Samples were centrifuged for 30 minutes at 21,000 g and 4°C, affording a two-phase partition, separated by insoluble protein residue. Polar metabolites and media aliquots were washed in methanol twice, then methoxymated by 20mg/mL methoxyamine (Sigma-Aldrich) in pyrimidine overnight and then trimethylsilyl derivatized in 99:1 BSTFA+TMS (Supelco

Analytical, Bellefonte PA) for 1 hour. Protein were extracted from the insoluble residue by homogenization in a 2% sodium dodecyl sulfate (SDS), 62.5mM Tris, and 1mM DTT, pH 6.8 buffer for protein determination using the Pierce BCA method (Thermo Fisher Scientific, Rockford, IL).

For *in vivo* metabolomic tracing experiments PDAC-265-GFP-luc cells were orthotopically injected into the pancreas of nude mice. Once liver metastasis onset was detected by IVIS, each mouse was given a bolus injection of $^{13}$C-U-glucose (20mg per 35g of body weight, 100ul of solution in saline). After 15 minutes an animal was culled and tumor and liver metastasis were dissected on ice and snap frozen. Tissue samples were then ground in liquid nitrogen to form a powder, which was then lyophilized. For metabolite extraction, ~10 mg of dry tissue (exact weights recorded) were extracted in 2ml Eppendorfs with Methanol:Chlorofom 1:2. First, 500μl of cold methanol containing 5uM $^{15}$N$^{13}$C-Valine, 10nmol scyllo-inositol and 10nmol of L-Norleucine as internal standards were added to the powder in 2ml tube and vortexed. Then 1200ul chloroform was added to the tube. After extensive vortexing, the tubes were sonicated (3x 8min, 4°C) and centrifugated (centrifugation for 20min, 4°C, 20,000g). The first supernatant was dried in speed-vac and 500 μl of 2:1 Methanol:water was added to the pellet (first methanol, vortex and then water). After extensive vortexing, the tubes were sonicated (3x 8min, 4°C) and centrifugated (20min, 4°C, 20000g). The supernatant 2 was added to tube of supernatant 1 and dried in speed-vac. Separation of polar and apolar phase was performed by addition of 3:3:1 Methanol:water:Chloroform (300:300:200 μl), vortexing and centrifugation for 30min, 4°C, 20000g). Polar phase was split to be run by LC-MS (80 μl).

Samples were analyzed on an Agilent 7890A GC equipped with a 5975C triple axis detector MSD (Agilent Technologies, Santa Clara CA). Metabolites were separated on an Agilent J&W 122-5532G DB-5ms capillary column (30m x 0.25mm, 0.25μm film thickness), in splitless mode. The injector and transfer line temperatures were 270 and 280°C, respectively. The flow rate of helium carrier gas was 0.7mL/min. The oven temperature was programmed to hold at 70°C for 2min, increased to 295°C at a 12.5°C/min ramp rate, increased from 295°C to 320°C at a 25°C/min ramp rate, and held at 320°C for 3min. The MS was operated in scan mode, after a 6min solvent delay, with a range of 50-565 m/z and a scan-rate of 2.8 scans per sec. Metabolites were identified by matching retention times and fragmentation patterns to commercially available standards. Metabolite peaks were integrated at each isotopologue m/z using MassHunter Workstation software (Agilent Technologies). Peak areas were quantified based on peak areas of known standards, using norleucine as an internal standard. Metabolites levels were normalized to protein content. Mass isotopologues were stripped of the contribution from natural abundance, based on the chemical formula of the derivatized fragment quantified. Percent enrichment for an isotopologue was calculated by dividing the corrected intensity by the sum of corrected intensities of all isotopologues for that metabolite.
